# Pdap1 protects mature B lymphocytes from stress-induced cell death and promotes antibody gene diversification

**DOI:** 10.1101/2020.01.30.917062

**Authors:** Verónica Delgado-Benito, Maria Berruezo-Llacuna, Robert Altwasser, Wiebke Winkler, Sandhya Balasubramanian, Devakumar Sundaravinayagam, Robin Graf, Ali Rahjouei, Madlen Driesner, Lisa Keller, Martin Janz, Altuna Akalin, Michela Di Virgilio

**Affiliations:** Laboratory of DNA Repair and Maintenance of Genome Stability, Max Delbrück Center for Molecular Medicine in the Helmholtz Association, Berlin 13125, Germany; Bioinformatics and Omics Data Science technology platform, Berlin Institute of Medical Systems Biology, Max Delbrück Center for Molecular Medicine in the Helmholtz Association, Berlin 13125, Germany; Laboratory of Immune Regulation and Cancer, Max Delbrück Center for Molecular Medicine in the Helmholtz Association, Berlin 13125, Germany; Laboratory of Biology of Malignant Lymphomas, Experimental and Clinical Research Center, Max Delbrück Center for Molecular Medicine in the Helmholtz Association and Charité, University Medicine, Berlin 13125, Germany

**Keywords:** Class switch recombination, somatic hypermutation, AID, mature B cells, Pdap1, integrated stress response

## Abstract

The establishment of protective humoral immunity is dependent on the ability of mature B cells to undergo antibody gene diversification while adjusting to the physiological stressors induced by activation with the antigen. Mature B cells diversify their antibody genes by class switch recombination (CSR) and somatic hypermutation (SHM), which are both dependent on efficient induction of activation-induced cytidine deaminase (AID). Here, we identified PDGFA-associated protein 1 (Pdap1) as an essential regulator of cellular homeostasis in mature B cells. Pdap1 deficiency leads to sustained expression of the integrated stress response (ISR) effector activating transcription factor 4 (Atf4) and induction of the ISR transcriptional program, increased cell death, and defective AID expression. As a consequence, loss of Pdap1 reduces germinal center B cell formation and impairs CSR and SHM. Thus, Pdap1 protects mature B cells against chronic ISR activation and ensures efficient antibody diversification by promoting their survival and optimal function.

## Introduction

The diversity of our immunoglobulin (Ig) gene repertoire is the result of antibody diversification reactions occurring at different stages of B lymphocytes development (Dudley et al., 2005; Methot and Di Noia, 2017). Developing B cells in the bone marrow randomly assemble different gene segments (known as variable (V), diversity (D) and joining (J) genes) at the *Ig* heavy (*Igh*) and light (*Igl*) chain loci *via* V(D)J recombination (Roth, 2014; Dudley et al., 2005). This process generates unique antibody gene receptors with the potential to collectively recognize a formidable number of antigens. Mature B cells further diversify their *Ig* genes in the periphery *via* somatic hypermutation (SHM) and class switch recombination (CSR) (Pavri and Nussenzweig, 2011; Methot and Di Noia, 2017). SHM introduces point mutations into the variable V(D)J region of the *Ig* genes to generate higher-affinity variants. CSR recombines the *Igh* constant (C) regions to replace the C portion of the IgM heavy chain with one of alternative isotypes (IgG, IgA and IgE), thus diversifying the Ig effector function. SHM and CSR are crucial to mount protective humoral responses, as evidenced by primary human immunodeficiency syndromes that are caused by defects in these reactions (Durandy et al., 2013).

SHM and CSR are both dependent on the B cell-specific enzyme activation-induced cytidine deaminase (AID) (Revy et al., 2000; Muramatsu et al., 2000). AID deaminates cytosine residues to uracil in single-stranded DNA stretches at the variable regions of both *Igh* and *Igl* loci during SHM, and within special recombining elements (switch (S) regions) of the *Igh* during CSR (Bransteitter et al., 2003; Chaudhuri et al., 2003; Petersen-Mahrt et al., 2002; Pham et al., 2003; Sohail et al., 2003; Ramiro et al., 2003; Dickerson et al., 2003; Matthews et al., 2014; Methot and Di Noia, 2017). The resulting U:G mismatches are differentially processed to generate either mutations in the variable regions or DNA breaks in the S regions (Peled et al., 2008; Methot and Di Noia, 2017; Matthews et al., 2014). AID expression is induced when resting mature (naïve) B cells are activated by the antigen and T-cell interactions (Zhou et al., 2003; Sayegh et al., 2003; Gonda et al., 2003; Cunningham et al., 2004; Dedeoglu et al., 2004; Muramatsu et al., 1999). Antigen stimulation reprograms naïve B cells to exit the quiescent state, expand their cellular biomass, and undergo a proliferative burst within transient and anatomically distinct structures in secondary lymphoid organs called germinal centers (GCs) (Cyster and Allen, 2019; Victora and Nussenzweig, 2012). AID expression peaks in GC B cells (Cattoretti et al., 2006; Crouch et al., 2007; Roco et al., 2019). The GC reaction represents the end stage of B cell development as GC B cells differentiate into memory B cells or long-lived plasma cells that secrete high affinity antibodies.

The integrated stress response (ISR) is a homeostatic program activated by a variety of physiological and pathological stresses to promote cellular recovery (Ron and Walter, 2007; Pakos-Zebrucka et al., 2016). These stresses include both cell intrinsic and extrinsic stimuli, such as endoplasmic reticulum (ER) stress, mitochondrial dysfunction, hypoxia, and amino acid deprivation (Quirós et al., 2017; Harding et al., 2003; Dever et al., 1992; Rzymski et al., 2010; Ye et al., 2010; Harding et al., 1999). All forms of stress converge into the phosphorylation of the alpha subunit of the eukaryotic translation initiation factor 2 (eIF2*α*) on serine 51 (Donnelly et al., 2013). This event causes a reduction in global protein synthesis while allowing the preferential translation of few selected genes including the ISR effector activating transcription factor 4 (Atf4) (Hinnebusch, 2000; Harding et al., 2000; Scheuner et al., 2001; Lu et al., 2004). Atf4 induces the transcriptional upregulation of stress-responsive genes and rewires cell metabolism towards the recovery of cellular homeostasis (Harding et al., 2003). The inhibition of general protein translation in the early stage of the ISR is followed by a later phase of translational recovery, which restores protein synthesis once the stress is resolved to support cell survival (Brostrom and Brostrom, 1998; Novoa et al., 2003; Kojima et al., 2003; Marciniak et al., 2004; Brostrom et al., 1989; Ma and Hendershot, 2003).

Although the ISR is an adaptive program meant to restore cellular homeostasis and to promote cell survival, under conditions of severe or prolonged stress, it induces cell death by activating pro-apoptotic pathways (Zou et al., 2008; Puthalakath et al., 2007; Teske et al., 2013; Wang et al., 2009; Ohoka et al., 2005; Gupta et al., 2012; Hiramatsu et al., 2014; Marciniak et al., 2004). Furthermore, the sustained protein synthesis during chronic stress causes proteotoxicity and leads to cell death (Han et al., 2013; Krokowski et al., 2013). Here, we identified PDGFA-associated protein 1 (Pdap1) as an essential regulator of mature B cell physiology that functions by countering chronic activation of the ISR. Pdap1 ablation in mature B cells causes sustained expression of Atf4, long-term induction of the ISR transcriptional program, and cell death. Furthermore, Pdap1 is essential for efficient induction of AID expression and physiological levels of CSR and SHM.

## Results

### Pdap1 is required for efficient CSR

To identify novel modulators of mature B cell physiology, we performed loss-of-CSR screens *via* somatic gene targeting by CRISPR-Cas9 in the B cell lymphoma line CH12. CH12 cells share several properties of resting B cells, and have been employed to study key aspects of late B cell differentiation, including B cell activation, antibody diversification by CSR, and immunoglobulin secretion (Arnold et al., 1983; Stockdale et al., 1987; Wiest et al., 1990; Corley et al., 1985; Bishop and Haughton, 1986; Nakamura et al., 1996; Ovnic and Corley, 1987; Fagone et al., 2007; Gass et al., 2002; Dufort et al., 2014; LoCascio et al., 1984; Locascio et al., 1984). Upon cytokine stimulation, CH12 cells express AID and undergo CSR to IgA with high efficiency (Nakamura et al., 1996). CRISPR-Cas9-mediated deletion of factors required for CSR reduces the efficiency of isotype switching in this cell line (Delgado-Benito et al., 2018). Among all tested candidates, targeting of Pdap1 resulted in a considerable reduction of CSR (Fig. 1, A and B). To confirm the results of the loss-of-CSR screen, we generated Pdap1-deficient CH12 clonal derivative cell lines, which included both indel knock-out (*Pdap1^−/−^*) and in-frame deletion mutant (*Pdap1^mut^*) clones (Fig. 1, C and D). In agreement with the CSR defect observed in bulk CH12 cultures, CSR was impaired in Pdap1-deficient clonal derivatives (Fig. 1E). We concluded that Pdap1 supports efficient CSR in CH12 cells.

**Figure 1.**
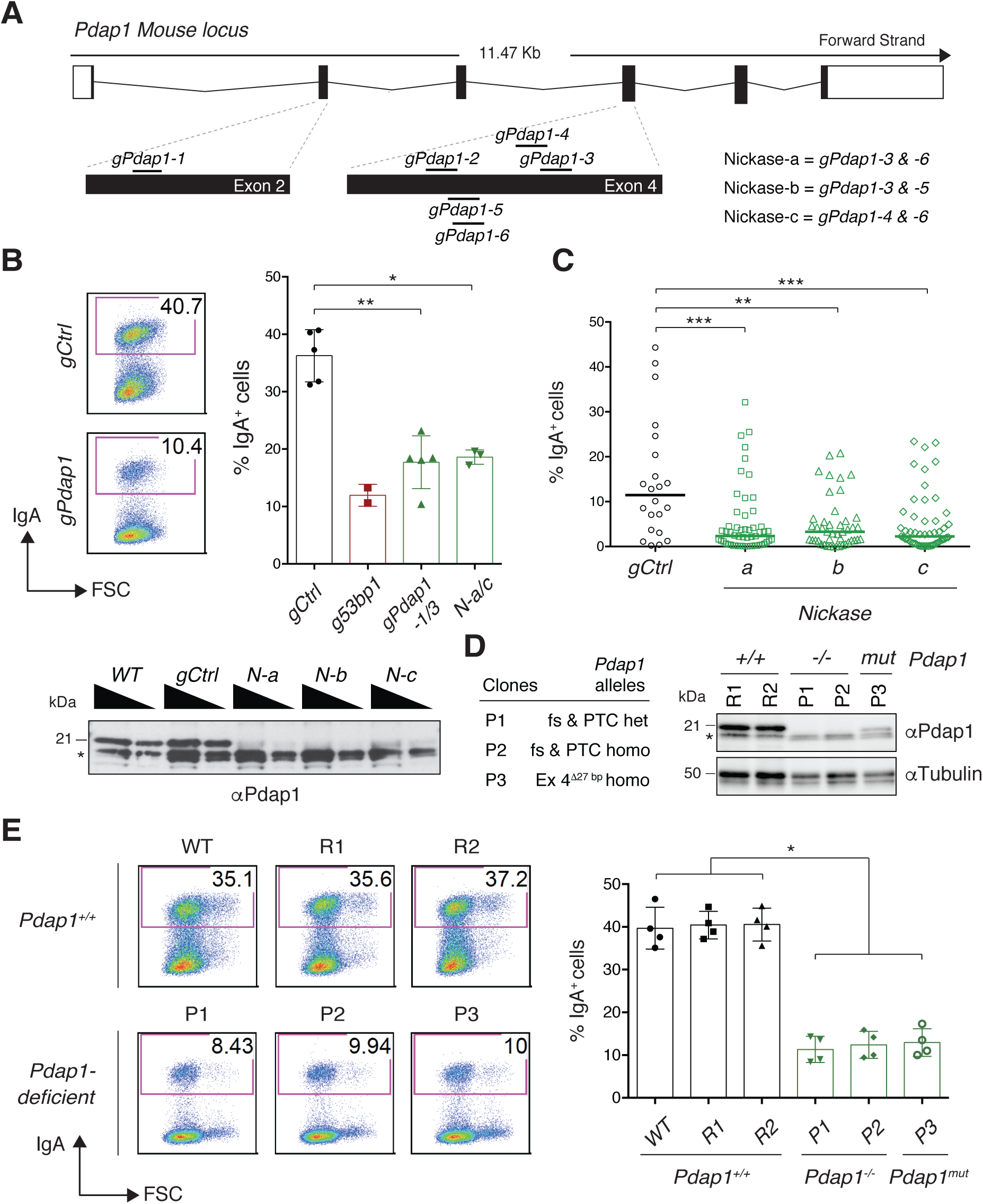
Loss of Pdap1 in CH12 cells impairs CSR. **A.** Scheme of murine *Pdap1* genomic locus and location of gRNAs used for gene targeting (scheme adapted from Ensembl Pdap1-201 ENSMUST00000031627.8). **B.** Top left: Representative flow cytometry plots measuring CSR to IgA in activated Cas9/*gPdap1*-nucleofected CH12 cells. Numbers in the plots refer to the percentage of switched cells (= IgA^+^ cells). Top right: Summary graph for three independent experiments employing *gPdap1-1/3* (individually / in pooled format) or *Nickases-a/c* (individually). Controls for gRNA-nucleofected CH12 were cells nucleofected either with empty vector or gRNAs against random sequences not present in the mouse genome (*gCtrl*). A previously described gRNA against the CSR factor 53BP1 (Delgado-Benito et al., 2018) was used as a positive control for loss-of-CSR function. Bottom: Western blot analysis of whole cell extracts from CH12 cultures nucleofected with *Nickases-a* to *-c*. Triangles indicate twofold dilution. Asterisk denotes aspecific bands used for internal normalization of protein levels. *N*: *Nickase*. **C.** Graph summarizing CSR efficiency of activated CH12 clonal cell lines derived by nucleofection of CH12 cultures with *Nickases-a/c* and single cell sorting. Each symbol in the graphs indicates a single cell clonal derivative. **D.** Left: Genomic scars of the targeted alleles in the selected Pdap1-deficient CH12 clonal derivatives. fs: frameshift; PTC: premature termination codon; Δ: bp deletion; het: heterozygous configuration (different indels causing fs and PTC at the two *Pdap1* alleles); homo: homozygous configuration. Right: Representative WB analysis of WT and Pdap1-deficient CH12 cell lines. R1 and R2 are WT clonal derivatives generated by targeting CH12 with random sequences not present in the mouse genome. *mut*: mutated. Asterisk denotes aspecific bands. **E.** Left: Representative flow cytometry plots measuring CSR to IgA in activated CH12 cell lines of the indicated genotypes. WT controls included both the parental CH12 cell line (WT) and the random clonal derivatives R1 and R2. Right: Summary graph for four independent experiments. Significance in panels B, C and E was calculated with the Mann–Whitney U test, and error bars represent SD. * = p ≤ 0.05; ** = p ≤ 0.01; *** = p ≤ 0.001.

Pdap1 is a 28 kDa phosphoprotein highly conserved in vertebrates. It was originally identified as a casein kinase II substrate and a weak interactor of platelet-derived growth factor (PDGF)-A (Shen et al., 1996; Fischer and Schubert, 2002). More recently, Pdap1 was described as an RNA-binding protein in several RNA-protein interactome studies (Castello et al., 2012, 2016; Trendel et al., 2019; Baltz et al., 2012; Iadevaia et al., 2020). However, the precise cellular function(s) of Pdap1 and its involvement in adaptive immunity are unknown.

To verify Pdap1 involvement in CSR *in vivo* and elucidate the underlying mechanism, we generated a mouse model bearing a conditional *Pdap1^F^* allele, and bred it to *Cd19^Cre/+^* mice to specifically ablate Pdap1 expression at the early stages of B cell differentiation (Rickert et al., 1997) (Fig. S1, A and B). B cell development was largely unaffected in *Pdap1^F/F^Cd19^Cre/+^* mice (Fig. S1, C and D), indicating that Pdap1 is dispensable for V(D)J recombination and early B cell differentiation. However, we observed a pronounced reduction in the number of splenic resting mature B cells (Fig. S1E). To confirm the intrinsic CSR defect caused by Pdap1-deficiency, we isolated resting splenocytes from *Pdap1^F/F^Cd19^Cre/+^* mice and monitored their capability to undergo CSR upon *in vitro* stimulation. We assessed CSR under conditions that induce switching to IgG1, IgG3, IgG2b, and IgA, and found that Pdap1-deficient B cells displayed reduced levels of CSR for all tested isotypes (Fig. 2, A to D). Western blot analysis confirmed the near complete abrogation of Pdap1 expression in these cells (Fig. 2E). Altogether, these data indicate that Pdap1 is required for efficient CSR in primary B cells irrespective of the stimulation condition.

**Figure 2.**
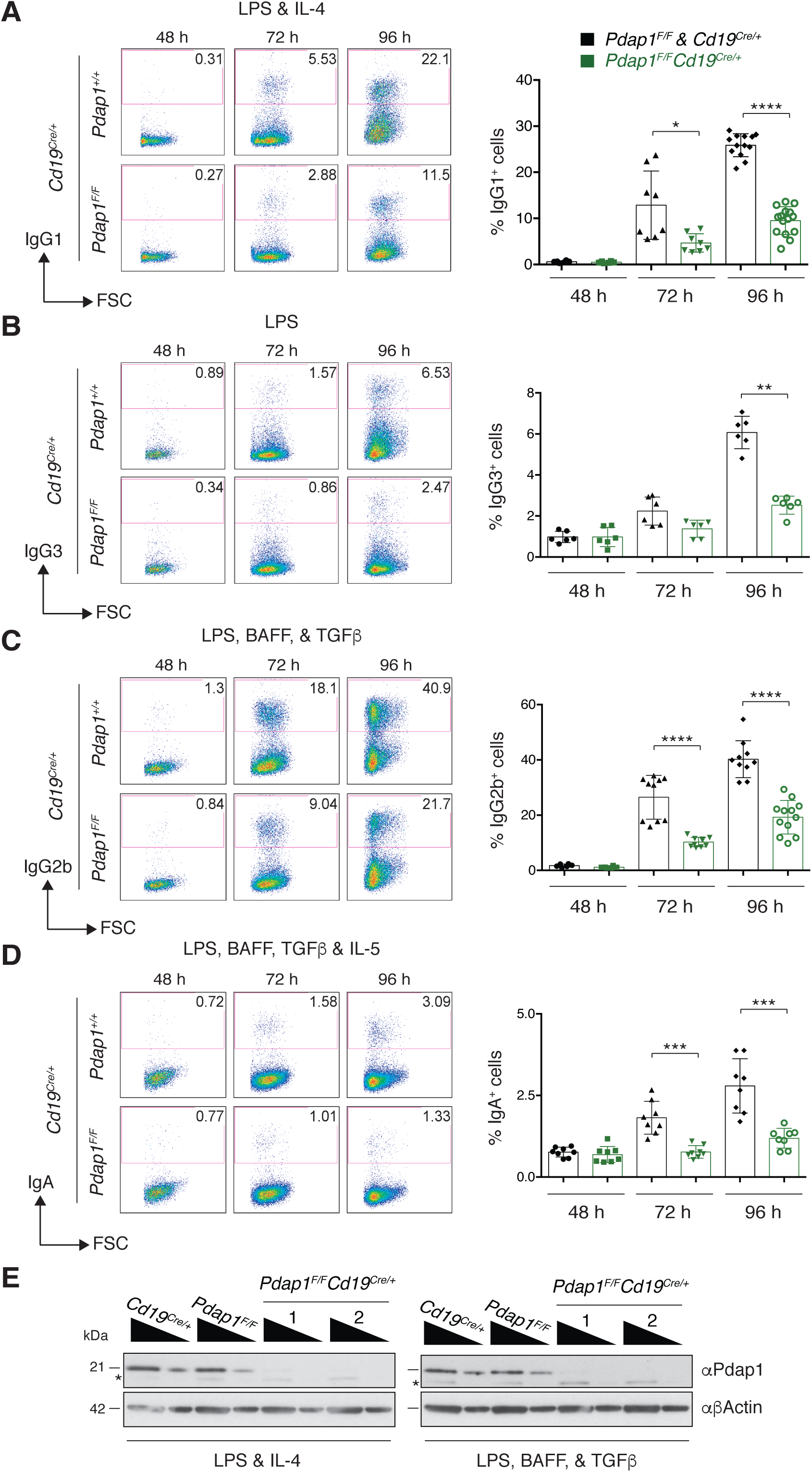
Pdap1 promotes CSR in primary B cells. **A-D.** Left: Representative flow cytometry plots measuring CSR to IgG1 (A), IgG3 (B), IgG2b (C), and IgA (D) in splenocytes activated with LPS and IL-4 (A), LPS (B), LPS, BAFF, and TGFβ (C), or LPS, BAFF, TGFβ, and IL-5 (D). Numbers in the plots refer to the percentage of switched cells (= IgG1^+^/IgG3^+^/IgG2b^+^/IgGA^+^ cells). Right: Summary graph for at least 6 mice per time point per genotype. **E.** Western blot analysis of whole cell extracts from B cells of the indicated genotypes 96 h after stimulation with LPS and IL-4 (left) or LPS, BAFF, and TGFβ (right). Numbers 1 and 2 indicate two different *Pdap1^F/F^Cd19^Cre/+^* mice. Triangles indicate threefold dilution. Asterisk denotes aspecific bands. Significance in panels A to D was calculated with the Mann–Whitney U test, and error bars represent SD. * = p ≤ 0.05; ** = p ≤ 0.01; *** = p ≤ 0.001; **** = p ≤ 0.0001. See also Figures S1 and S2.

### Pdap1 supports physiological levels of AID expression

CSR is dependent on cell proliferation (Hasbold et al., 1998; Hodgkin et al., 1996; Deenick et al., 1999; Hasbold et al., 1999). To determine if the reduced CSR efficiency of Pdap1-deficient B cells is caused by an underlying defect in cell proliferation, we monitored the proliferation capabilities of splenocytes cultures by cell tracking dye dilution. The CellTrace Violet dilution profiles of *Pdap1^F/F^Cd19^Cre/+^* B cells were indistinguishable from the control counterparts under all stimulation conditions (Fig. S2, A to D). Furthermore, class switching was reduced in Pdap1-deficient B cells independently of the number of cell divisions (Fig. S2E). We concluded that Pdap1 is dispensable for B cell proliferation, and that the CSR defect of *Pdap1^F/F^Cd19^Cre/+^* B cells is not due to reduced proliferation capabilities.

The efficiency of CSR is directly linked to the levels of AID expression (Dorsett et al., 2008; Takizawa et al., 2008; Teng et al., 2008). Furthermore, AID targeting is dependent on non-coding transcription across the S regions (germline transcription, GLT), which exposes single-stranded DNA stretches that are the substrate of AID-mediated deamination (Chaudhuri et al., 2003; Dickerson et al., 2003; Ramiro et al., 2003). Therefore, we monitored *Aicda* mRNA and GLT levels in activated B cells by quantitative reverse transcription PCR (RT-qPCR) analysis. We found that *Aicda* transcript levels were consistently reduced in *Pdap1^F/F^Cd19^Cre/+^* B cells compared to controls across all stimulation conditions (Fig. 3A). Transcription of donor Sμ region was not affected by Pdap1 deletion (Fig. 3B), whereas acceptor S region transcription exhibited a varied phenotype, with reduced levels of GLT*γ*1 and GLT*α*, and minimally affected or unaltered expression for GLT*γ*3 and GLT*γ*2b, respectively (Fig. 3C). Analogously, analysis of *Aicda* and germline transcripts in Pdap1-deficient CH12 cell lines showed reduced *Aicda* mRNA levels but unaffected Sμ and S*α* region transcription (Fig. S3, A to C). In agreement with the reduction of *Aicda* mRNA, Pdap1-deficient splenocytes expressed lower levels of AID protein upon activation compared to control cells (Fig. 3D).

**Figure 3.**
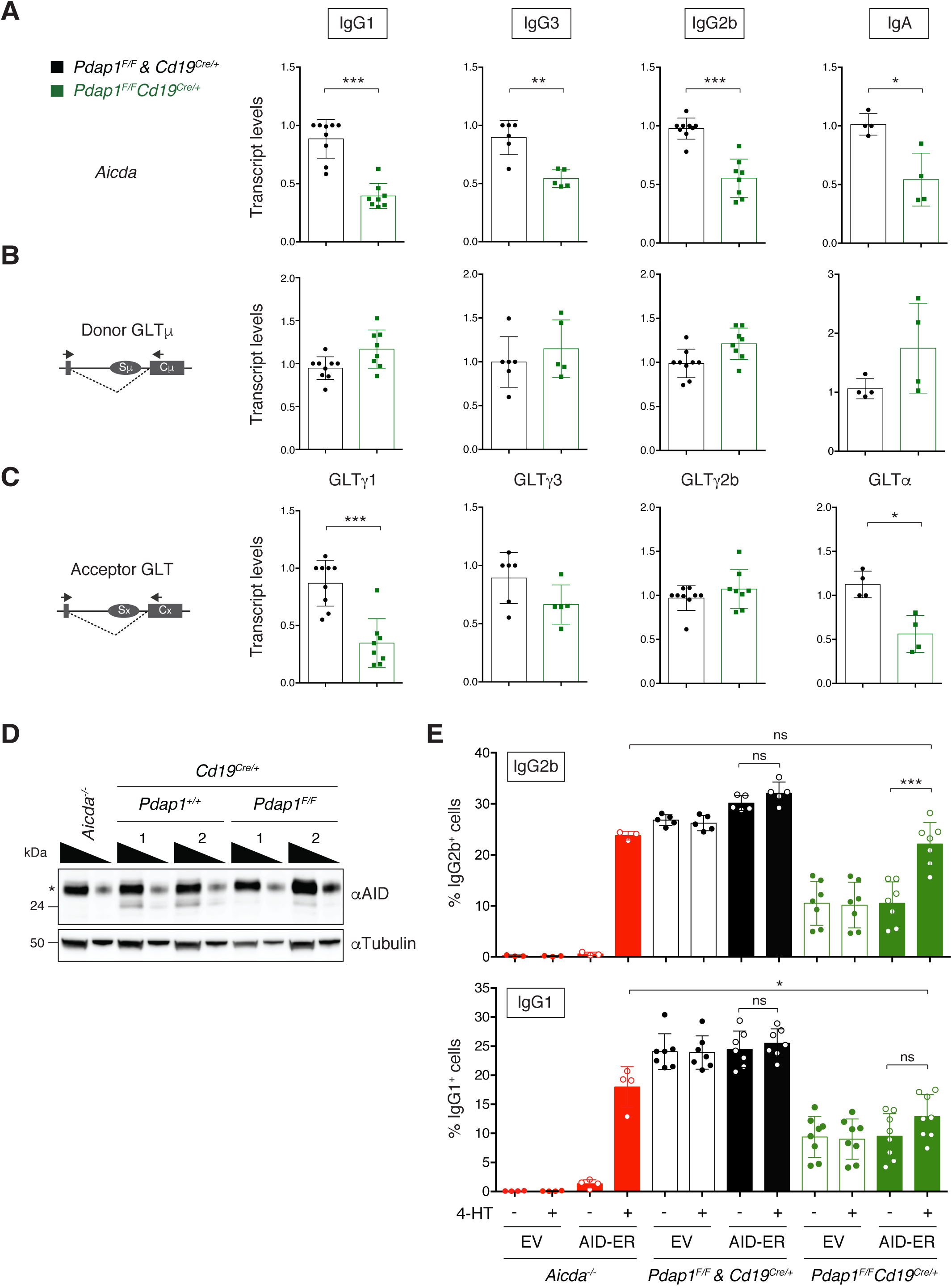
Pdap1 supports physiological levels of AID expression. **A-C.** qPCR analysis for *Aicda* mRNA (A), Igμ (B), and Ig*γ*1 / Ig*γ*3 / Ig*γ*2b / Ig*α* (C) GLT levels in B cells activated to undergo CSR to the corresponding isotypes. The schematic representations on the left in panels B and C indicate the location of primers employed to analyze the germline transcripts. Graphs summarize 4 to 8 mice per genotype per stimulation condition. One control mouse (*Cd19^Cre/+^* or *Pdap1^F/F^*) within each experiment was assigned an arbitrary value of 1. **D.** Representative WB analysis of splenocytes of the indicated genotypes 48 h after stimulation with LPS and IL-4. Numbers 1 and 2 indicate two different mice per genotype. Triangles indicate threefold dilution. Asterisk denotes aspecific bands. **E.** Summary graphs showing CSR to IgG2b (top) and IgG1 (bottom) following transduction of splenocytes of the indicated genotypes with either empty vector (EV) or an AID-ER-expressing retroviral construct. 4-HT: 4-hydroxytamoxifen. Graphs summarize 3-8 mice per genotype analysed in three (IgG2b) or four (IgG1) independent experiments. Significance in panels A, C and E was calculated with the Mann–Whitney U test. Error bars represent SD. ns: not significant; * = p ≤ 0.05; ** = p ≤ 0.01; *** = p ≤ 0.001. See also Figures S3 and S4.

To confirm that the reduced CSR efficiency of *Pdap1^F/F^Cd19^Cre/+^* B cells is due to lower levels of AID expression, we assessed whether overexpression of AID in these cells could rescue the CSR defect. To this end, we transduced *Cd19^Cre/+^* and *Pdap1^F/F^Cd19^Cre/+^* splenocytes with a construct expressing AID fused to the hormone-binding domain of the modified estrogen receptor (AID-ER). Provision of 4-hydroxytamoxifen (4-HT) induces the translocation of the cytoplasm-accumulated AID-ER fusion protein into the nucleus and initiates CSR. We found that 4-HT addition rescued the CSR defect of *Pdap1^F/F^Cd19^Cre/+^* cells to levels equivalent to AID-ER-reconstituted *Aicda^−/−^* splenocytes in cultures stimulated to undergo CSR to IgG2b but not in cells stimulated to undergo CSR to IgG1 (Fig. 3E). This result is in agreement with the additional defect in acceptor GLT expression exhibited by *Pdap1^F/F^Cd19^Cre/+^* B cells switching to IgG1 (Fig. 3C). We concluded that Pdap1-deficiency impairs AID expression, and as a consequence, CSR.

Pdap1 has been recently identified in RNA-protein capture experiments as an RNA binding protein (Castello et al., 2012, 2016; Trendel et al., 2019; Baltz et al., 2012; Iadevaia et al., 2020). Therefore, we considered the possibility that Pdap1 might contribute to the post-transcriptional regulation of AID expression. To test this hypothesis, we treated *Cd19^Cre/+^* and *Pdap1^F/F^Cd19^Cre/+^* splenocytes cultures with Actinomycin D to inhibit the *de novo* transcription of AID, and measured *Aicda* mRNA decay by qRT-PCR. We found that depletion of Pdap1 did not affect the kinetics of *Aicda* mRNA degradation (Fig. S4, A and B). This result indicates that Pdap1 is dispensable for *Aicda* transcript stability, decapping and deadenylation since a defect in any of these processes would result in accelerated decay. Next, we monitored *Aicda* transcript splicing. Several splice variants for AID have been identified, with the full-length transcript being the only splice isoform encoding a functional AID protein (van Maldegem et al., 2009; Sala et al., 2015; van Maldegem et al., 2010). To assess whether Pdap1 modulates *Aicda* splicing, we performed RNA next-generation sequencing (RNA-Seq) of activated B cells induced to undergo CSR to either IgG1 or IgG2b, and analysed the pattern of *Aicda* exon usage. We found no difference in the profile of *Aicda* exon usage between *Cd19^Cre/+^* and *Pdap1^F/F^Cd19^Cre/+^* B cells (Fig. S4C). Furthermore, primary *Aicda* transcript levels in *Pdap1^F/F^Cd19^Cre/+^* splenocytes were reduced compared to *Cd19^Cre/+^* cells to the same extent observed for the full-length spliced product (Fig. S4D and 3A). We concluded that ablation of Pdap1 does not affect *Aicda* mRNA splicing.

Altogether, these findings indicate that Pdap1 supports physiological levels of AID expression, and that the CSR defect of Pdap1-deficient B cells is caused, at least in part, by impaired induction of AID expression following activation rather than a defect in *Aicda* post-transcriptional regulation.

### Pdap1 deficiency in B cells induces the Atf4 stress response transcriptional program

To define the mechanism responsible for the defective induction of AID expression in Pdap1-deficient cells, we compared the transcriptome profiles of *Cd19^Cre/+^* and *Pdap1^F/F^Cd19^Cre/+^* splenocytes stimulated with either LPS and IL-4, or LPS, BAFF, and TGFβ (Table S1). The use of two stimulation conditions allowed us to zoom in common differentially-regulated pathways, and added a temporal dimension to the experiment since LPS-BAFF-TGFβ-stimulated splenocytes proliferate faster than LPS-IL-4-activated ones (Fig. S2F). To identify differentially expressed genes, we set the significance level of false discovery rate (FDR) to < 0.05 (Fig. 4A). The number of differentially regulated genes was higher in LPS-IL-4-stimulated cultures than in LPS-BAFF-TGFβ-activated cells (1227 *versus* 173). Furthermore, the number of genes upregulated in *Pdap1^F/F^Cd19^Cre/+^* was considerably higher than the downregulated ones in LPS-BAFF-TGFβ-stimulated cultures (124 up- *versus* 49 downregulated), but was evenly distributed among the two categories in the LPS-IL-4 stimulation condition (561 up- *versus* 666 downregulated).

**Figure 4.**
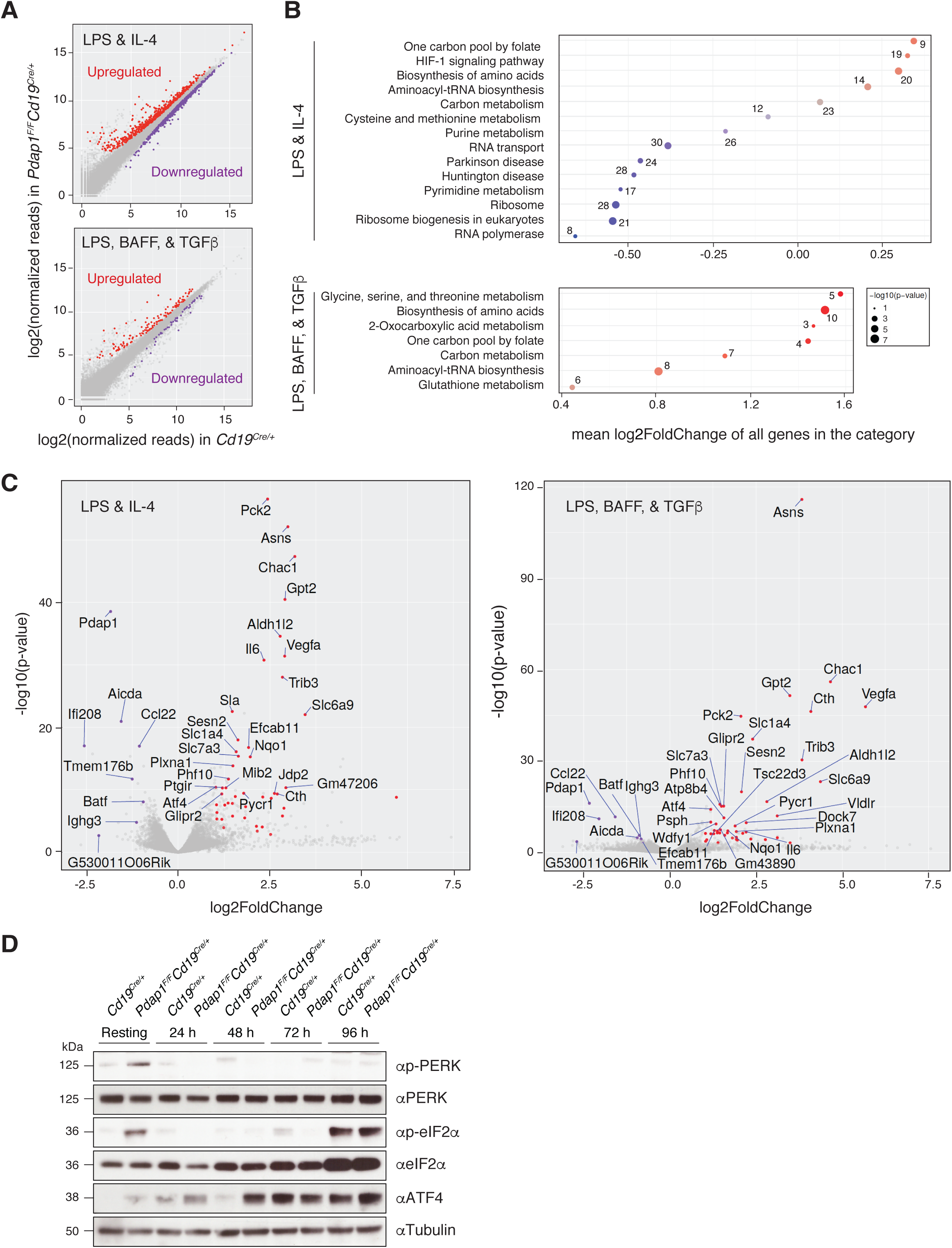
Pdap1-deficient B cells activate the Atf4 stress response transcriptional program. **A.** Scatterplots of gene expression in *Pdap1^F/F^Cd19^Cre/+^ versus* control (*Cd19^Cre/+^*) mature B cells stimulated with LPS and IL-4 (top) or LPS, BAFF, and TGFβ (bottom). Genes with an adjusted p-value (FDR) < 0.05 that are up- or downregulated in *Pdap1^F/F^Cd19^Cre/+^* cells are highlighted in red or purple, respectively. Data summarizes results from 3 mice per genotype per stimulation condition, and is presented as log2(RPM) (reads per gene per million mapped sequence reads) values. **B.** Pathway enrichment analysis (KEGG pathways) of the differentially regulated genes (FDR < 0.05) from panel A. The number of differentially regulated genes in each category is indicated. **C.** Volcano plots displaying differentially expressed genes between control and Pdap1-deficient splenocytes activated with LPS and IL-4 (right) or LPS, BAFF, and TGFβ (left). The red and purple dots represent transcripts up- and downregulated in *Pdap1^F/F^Cd19^Cre/+^* cells, respectively, with FDR < 0.05 and expression fold change > 2 (upregulated) or 1.7 (downregulated) in both stimulation conditions. The names of the downregulated and the 30 most significantly upregulated genes within each stimulation condition are indicated in each graph. The fold change threshold for downregulated genes was set to 1.7 to include genes yielding a biologically relevant effect even with less pronounced variations in expression levels (e.g. *Aicda*, haploinsufficient gene). **D.** Representative WB analysis of splenocytes of the indicated genotypes before (resting B cells) and after 24 to 96 h activation with LPS and IL-4. Data is representative of 2 mice per genotype. See also Figure S5 and Table S1.

Pathway enrichment analysis of the downregulated genes in *Pdap1^F/F^Cd19^Cre/+^* B cells in the LPS-IL-4 data set identified the protein translation-related categories of “Ribosome”, “Ribosome biogenesis in eukaryotes”, and “RNA transport” as the most significant categories (Fig. 4B). Furthermore, analysis of individual genes showed that the most significantly downregulated ones under both stimulations were factors essential for CSR and/or genes highly expressed or induced in activated B cells (e.g. *Aicda*, *Batf*, *Ccl22, Ighg3, Tmem176b*) (Fig. 4C and Table S1). In contrast, the top upregulated categories in both datasets were related to metabolic pathways of amino acid and aminoacyl-tRNA biosynthesis (Fig. 4B). Among the most significantly upregulated genes, we found key enzymes for the biosynthesis of asparagine (*Asns*), cysteine (*Cth*), and serine (*Psat1*, *Psph*), glutathione (GSH) metabolism (*Chac1*), amino acid transporters (*Slc1a4*, *Slc6a9*), and several other metabolic and cytoprotective genes (Fig. 4C, S5, and Table S1).

These data indicate that, in addition to the inhibition of key factors of the CSR program, Pdap1 deficiency in activated B cells is accompanied by transcriptional changes that reflect initial repression of global protein synthesis, up-regulation of cellular stress response genes, and metabolic rewiring to aid translational recovery. These changes represent key aspects of the transcriptional program of the ISR (Ron and Walter, 2007; Pakos-Zebrucka et al., 2016). To confirm the activation of the ISR in Pdap1-deficient B cells, we monitored the kinetics of eIF2*α* phosphorylation and Atf4 expression in resting and *in vitro*-activated splenocytes. The eIF2*α*-Atf4 pathway is normally suppressed in splenic B cells, and is induced only at later time points after activation, when splenocyte cultures exhibit a general fitness decline (Fig. 4D, control cells, and (Zhu et al., 2019)). In contrast to control cells, resting B cells from *Pdap1^F/F^Cd19^Cre/+^* spleens displayed a marked phosphorylation of eIF2*α* and detectable levels of Atf4 protein, thus indicating a baseline activation of the ISR (Fig 4D). Following activation, eIF2*α* phosphorylation was no longer detectable in both control and Pdap1-deficient cells (Fig. 4D). The autophosphorylation-dependent activation of the ER stress-activated eIF2*α* kinase PKR-like ER kinase (Perk) mirrored the phosphorylation profile of eIF2*α* (Fig. 4D). In contrast, Atf4 protein levels in *Pdap1^F/F^Cd19^Cre/+^* cells were more pronounced at 24 h after activation and increased steadily over time (Fig. 4D). We concluded that Pdap1 deficiency results in upregulation of the ISR core component Atf4.

Altogether, these data indicate that Pdap1 ablation in mature B cells induces Atf4-dependent expression of stress response genes and downregulation of the CSR program.

### Pdap1 supports survival of mature B cells

The ISR is an adaptive pathway meant to restore cellular homeostasis following cell intrinsic and extrinsic stresses (Harding et al., 2003, 2000; Brostrom et al., 1996; Dever et al., 1992; Ron and Walter, 2007; Pakos-Zebrucka et al., 2016; Harding et al., 1999). However, when the stress is severe in either intensity or duration and overwhelms the response adaptive capacity, the ISR activates the apoptotic cell death program (Pakos-Zebrucka et al., 2016). Furthermore, downregulation studies have implicated Pdap1 in cell survival and apoptosis resistance of cancer cell lines (Sharma et al., 2016; Weston et al., 2018). Therefore, we tested whether activation of the stress response in the absence of Pdap1 would induce apoptosis in B cells. To do so, we monitored caspase activation in splenocytes by CaspGLOW staining. Caspase activity increased after stimulation and was higher in Pdap1-deficient cells compared to controls at 24 and 48 h post-activation (Fig. 5A). Accordingly, despite the unaffected proliferation capability (Fig. S2), the number of live cells in *Pdap1^F/F^Cd19^Cre/+^* splenocytes cultures was reduced 48 and 72 h after activation (Fig. 5B). Finally, we found that a large proportion of the *Pdap1^F/F^Cd19^Cre/+^* B cell cultures displayed reduced mitochondrial membrane potential at early time points after activation, which is indicative of increased mitochondrial membrane depolarization (Fig. 5C).

**Figure 5.**
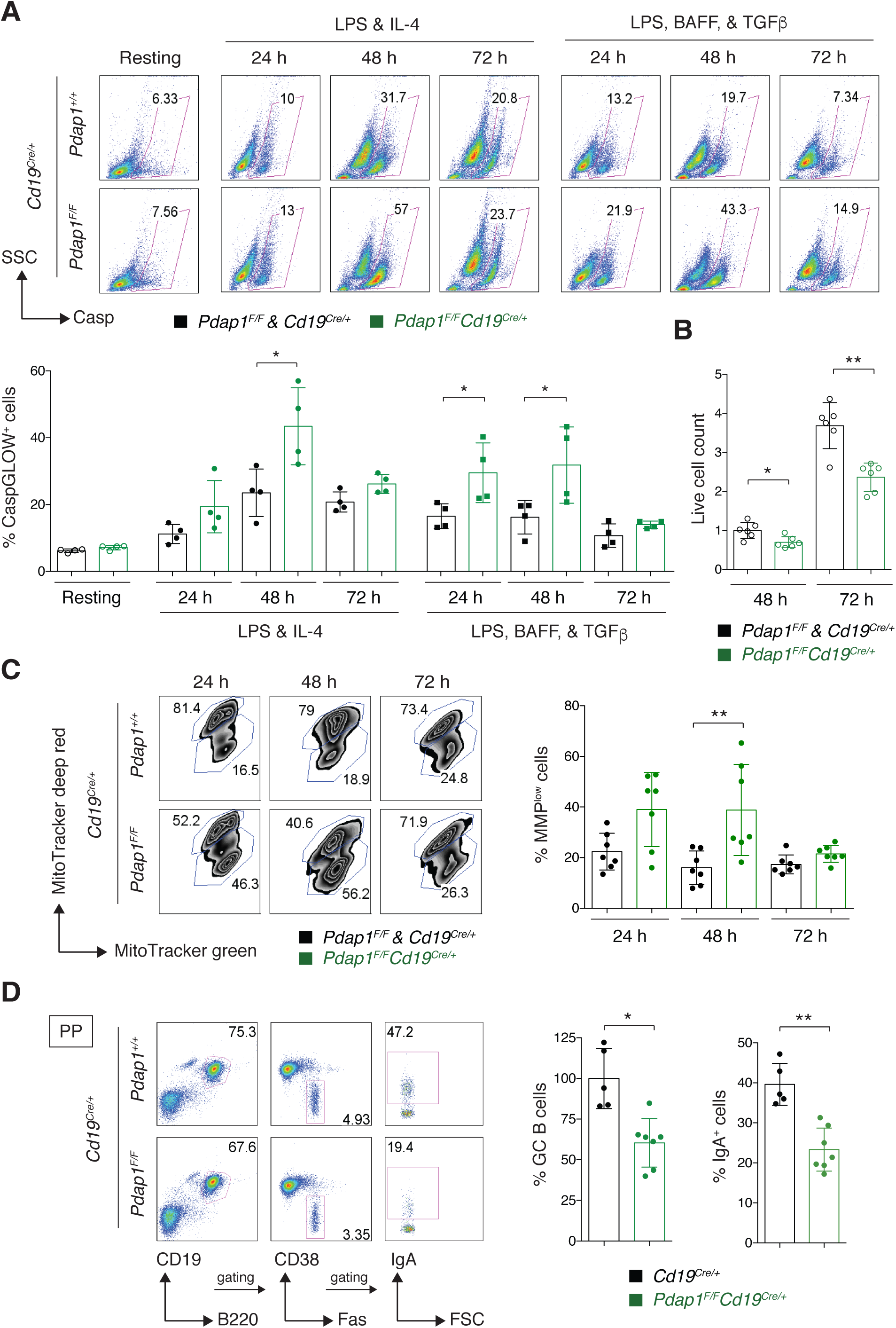
Pdap1 supports survival of mature B cells. **A.** Top: Representative flow cytometry plots measuring percentage of active caspases^+^ cells in resting and activated (LPS-IL-4, or LPS-BAFF-TGFβ, 24 to 72 h) splenocytes of the indicated genotypes. Bottom: Summary graph for 4 mice per genotype. **B.** Live cell count of splenocytes of the indicated genotypes activated with LPS and IL-4. Resting B cells were counted and seeded at same density on day 0. Data summarizes three independent experiments for a total of 6 mice per genotype, and is normalized within each experiment to the average of live cell count for control cultures at 48 h, which was set to 1. **C.** Left: Representative flow cytometry plots measuring mitochondrial membrane potential (MMP) and mass by MitoTracker deep red and green staining, respectively, in splenocyte cultures of the indicated genotype 24, 48, and 72 h after activation with LPS and IL-4. Right: Summary graph for 7 mice per genotype. **D.** Left: Representative flow cytometry plots measuring percentage of germinal center (GC) B cells and IgA^+^ GC cells in Peyer’s patches (PP) of unimmunized mice. Right: Summary graphs for at least 5 mice per genotype. Significance in panels A, B, C, and D was calculated with the Mann–Whitney U test, and error bars represent SD. * = p ≤ 0.05; ** = p ≤ 0.01.

Next, we tested whether Pdap1 deletion would affect the survival of activated B cells *in vivo*. To this end, we measured the percentage of GC B cells in Peyer’s patches of *Cd19^Cre/+^* and *Pdap1^F/F^Cd19^Cre/+^* mice. Peyer’s patches are specialized secondary lymphoid tissues that line the wall of the small intestine. Because of their chronic exposure to an enormous variety of food- and microbiome-derived antigens, Peyer’s patches display continual germinal center activity, and are key to the induction of mucosal IgA antibody responses (Reboldi and Cyster, 2016). We found that the percentage of GC B cells was reduced in Peyer’s patches of *Pdap1^F/F^Cd19^Cre/+^* mice compared to the control group (Fig. 5D). In agreement with the CSR defect exhibited by *Pdap1^F/F^Cd19^Cre/+^* splenic B cells following activation *in vitro* (Fig. 2), the percentage of switched IgA GC B cells was considerably decreased in *Pdap1^F/F^Cd19^Cre/+^* mice (Fig. 5D).

Altogether, these data suggest that Pdap1 is required to support survival of activated B cells.

### Pdap1 is dispensable for plasma cell differentiation

Plasma cell differentiation represents the terminal phase of B cell development, and is regulated by a transcriptional program that represses B cell identity while promoting the expression of plasma cell signature genes (Nutt et al., 2015; Shi et al., 2015; Minnich et al., 2016). This process is accompanied by massive expansion of the ER and the upregulation of molecular chaperone and folding enzyme expression. The expansion of the secretory network is essential to accommodate the demands of increased immunoglobulin synthesis and secretion, and is dependent on the activation of the unfolded protein response (UPR) (Zhang et al., 2005; Gass et al., 2002, 2008; Iwakoshi et al., 2003; Van Anken et al., 2003; Shaffer et al., 2004; Taubenheim et al., 2012; Todd et al., 2009). The UPR is a ubiquitous signaling network that senses and responds to the accumulation of misfolded proteins in the ER (Ron and Walter, 2007; Walter and Ron, 2011; Schroder and Kaufman, 2005). This response is mediated by three ER resident proteins, which in addition to Perk, comprise the inositol-requiring protein kinase/endoribonuclease-1 (Ire1) and the activating transcription factor 6 (Atf6) (Tirasophon et al., 1998; Wang et al., 1998; Haze et al., 1999, 2001). The UPR ultimately re-establishes protein homeostasis by integrating Perk-eIF2*α*-dependent attenuation of global protein synthesis with Ire1- and Atf6-mediated transcription of factors and enzymes that increase the protein folding and degradation capabilities of the ER.

The Ire1 and Atf6 branches of the UPR are activated during normal differentiation of naïve B cells into plasma cells, and the Ire1 arm is essential for the expansion of their secretory network (Zhang et al., 2005; Gass et al., 2002; Iwakoshi et al., 2003; Van Anken et al., 2003; Gass et al., 2008; Aragon et al., 2012). In contrast, the Perk arm of the UPR is dispensable for plasma cell development (Zhang et al., 2005; Gass et al., 2008). Furthermore, this pathway is suppressed during differentiation of naïve B cells into plasma cells (Ma et al., 2010; Gass et al., 2008; Zhang et al., 2005), and unrestricted Perk-eIF2*α*-Atf4 signaling in activated B cells blocks the formation of plasma cells (Zhu et al., 2019). Given the upregulated expression of Atf4 in Pdap1-deficient B cells, we considered the possibility that plasma cell development might be impaired in the absence of Pdap1. To this end, we analyzed the plasma cell compartment of *Pdap1^F/F^Cd19^Cre/+^* mice. We found that the percentage and number of plasma cells (CD138^+^TACI^+^) in bone marrow and spleen of unimmunized mice was similar between control and *Pdap1^F/F^Cd19^Cre/+^* mice (Fig. 6A).

**Figure 6.**
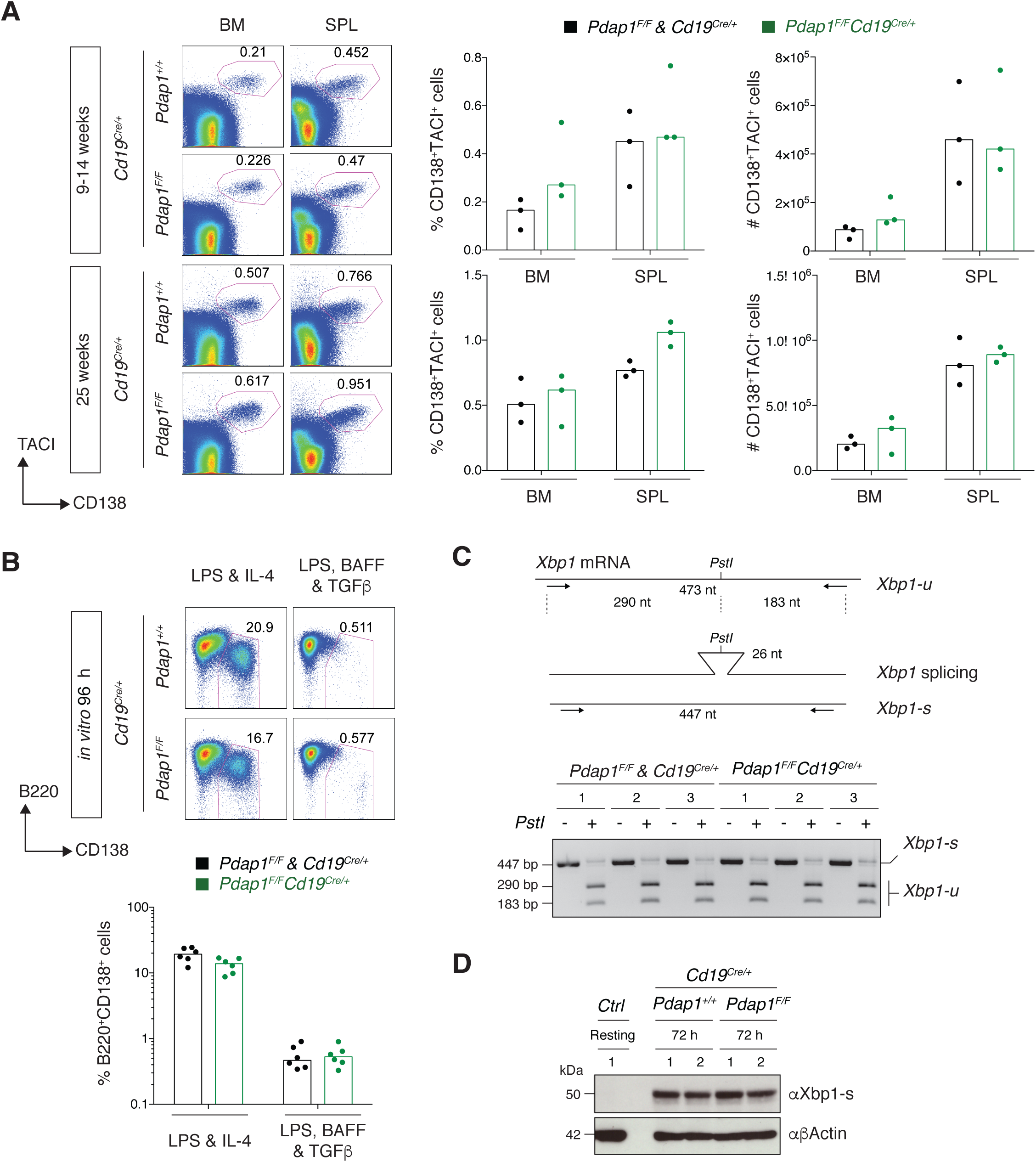
Pdap1 is dispensable for plasma cell differentiation. **A.** Left: Representative flow cytometry plots showing percentage of plasma cells (CD138^+^TACI^+^) in bone marrow (BM) and spleen (SPL) of unimmunized mice of the indicated genotypes and age. Right: Summary graphs showing percentage and number of plasma cells for 3 mice per genotype and age group. **B.** Top: Representative flow cytometry plots measuring percentage of plasmablasts (B220^int^CD138^+^) in splenocyte cultures of the indicated genotypes 96 h after activation with LPS and IL-4, or LPS, BAFF, and TGFβ. Bottom: Summary graph for 6 mice per genotype. **C.** Top: Schematic representation of *Xbp1* mRNA RT-PCR and digestion products. Bottom: Electrophoretic analysis of *Xbp1* splicing in splenocytes cultures of the indicated genotypes 48 h after activation with LPS and IL-4. The DNA bands of 447 bp for the intact spliced (Xbp1-s) transcript RT-PCR product, and the unspliced (Xbp1-u) transcript digestion products of 290 bp and 183 bp are indicated. Results for 3 mice per genotype are shown. **D.** Representative WB analysis of splenocytes of the indicated genotypes 72 h after activation with LPS and IL-4. Resting B cells from a *WT* control mouse (*Ctrl*) were analysed in parallel to show the expected undetectable levels of Xbp1-s before activation. Numbers 1 and 2 indicate two different mice per genotype. No difference among groups was significant for graphs in panels A and B (Mann–Whitney U test).

Activation of naïve B cells with specific combinations of stimuli *in vitro* induces differentiation into plasmablasts. Plasmablasts are proliferating antibody-secreting cells whose differentiation is driven by the same transcriptional program responsible for plasma cell development (Nutt et al., 2015). Therefore, we tested the formation of plasmablasts (B220^+^CD138^+^) *in vitro* under the same stimulation conditions employed for our previous analyses. We observed no significant difference in the percentage of plasmablasts between controls and *Pdap1^F/F^Cd19^Cre/+^* cultures (Fig. 6B). Accordingly, we did not detect any apparent change in the activation of the plasmablast and plasma cell gene signature between the two genotypes in the RNA-Seq analysis (Table S1). We concluded that Atf4 upregulation in activated *Pdap1^F/F^Cd19^Cre/+^* B cells does not inhibit the generation of plasmablasts and plasma cells, and that Pdap1 is not required for the B cell developmental program controlling plasmablast and plasma cell formation and identity.

The ER expansion and increase in protein processing and folding capabilities of plasma cells are regulated by the transcription factor Xbp1-s (spliced Xbp1) (Shaffer et al., 2004). Xbp1-deficiency in B cells does not interfere with the development of plasma cells, but impairs immunoglobulin secretion because of defective expansion of the secretory pathway (Hu et al., 2009; Todd et al., 2009; Taubenheim et al., 2012; Shaffer et al., 2004). Xbp1-s is expressed as a result of an unconventional splicing event mediated by Ire1 that removes a 26 nt segment from *Xbp1* mRNA (unspliced *Xbp1* transcript, *Xbp1-u*), thus changing the transcript reading frame in the resulting spliced *Xbp1* transcript (*Xbp1-s*) (Fig. 6C and (Yoshida et al., 2001)). Xbp1-s translocates into the nucleus and activates the transcription of genes responsible for the expansion of the secretory pathway (Shaffer et al., 2004; Tellier et al., 2016). We found no difference in *Xbp1* mRNA splicing and Xbp1-s expression levels between control and *Pdap1^F/F^Cd19^Cre/+^* splenocytes cultures (Fig. 6, C and D). These results suggest that the transcriptional program controlling ER network expansion in plasma cells is not affected by Pdap1 deletion.

We concluded that Pdap1 is dispensable for the differentiation programs that establish plasma cell identity and function.

### Pdap1 is required for efficient SHM of *Ig* loci

*Aicda* is an haploinsufficient gene, and *Aicda^+/-^* mice exhibit defects in both antibody diversification reactions initiated by AID, CSR and SHM (McBride et al., 2008; Sernández et al., 2008; Takizawa et al., 2008). Given the reduced levels of AID expressed by *Pdap1^F/F^Cd19^Cre/+^* B cells following activation, we considered the possibility that Pdap1 depletion might affect also SHM. To test this hypothesis, we sequenced the intronic regions downstream of the J_H_4 and J_K_5 elements in GC B cells sorted from Peyer’s patches of aged *Pdap1^F/F^Cd19^Cre/+^* mice (Jolly et al., 1997). Because of their chronic exposure to microbial antigens, GC B cells in Peyer’s patches accumulate a high number of mutations over time (González-Fernández et al., 1994). Accordingly, we found that GC B cells from control mice displayed a high mutation frequency at both J_H_4 and J_K_5 intronic regions (Fig. 7, A to D). The vast majority of sequences contained mutations (86%, 79/92 for J_H_4, and 83%, 104/125 for J_K_5), and a considerable portion of B cells was heavily mutated (Fig. 7, E and F). In contrast, *Pdap1^F/F^Cd19^Cre/+^* GC B cells exhibited a significantly lower mutation frequency at both J_H_4 and J_K_5 introns (Fig. 7, A to D). The distribution of mutations per sequence was skewed towards an increase in the proportion of sequences bearing no or less than 5 mutations (50%, 54/107 for J_H_4, and 65%, 83/127 for J_K_5), whereas the number of highly mutated clones was concomitantly reduced (Fig. 7, E and F). In agreement with a role of Pdap1 in promoting AID expression rather than the processing of AID-induced lesions, the profile of mutations was not affected by Pdap1 deletion (Fig. 7, G and H).

**Figure 7.**
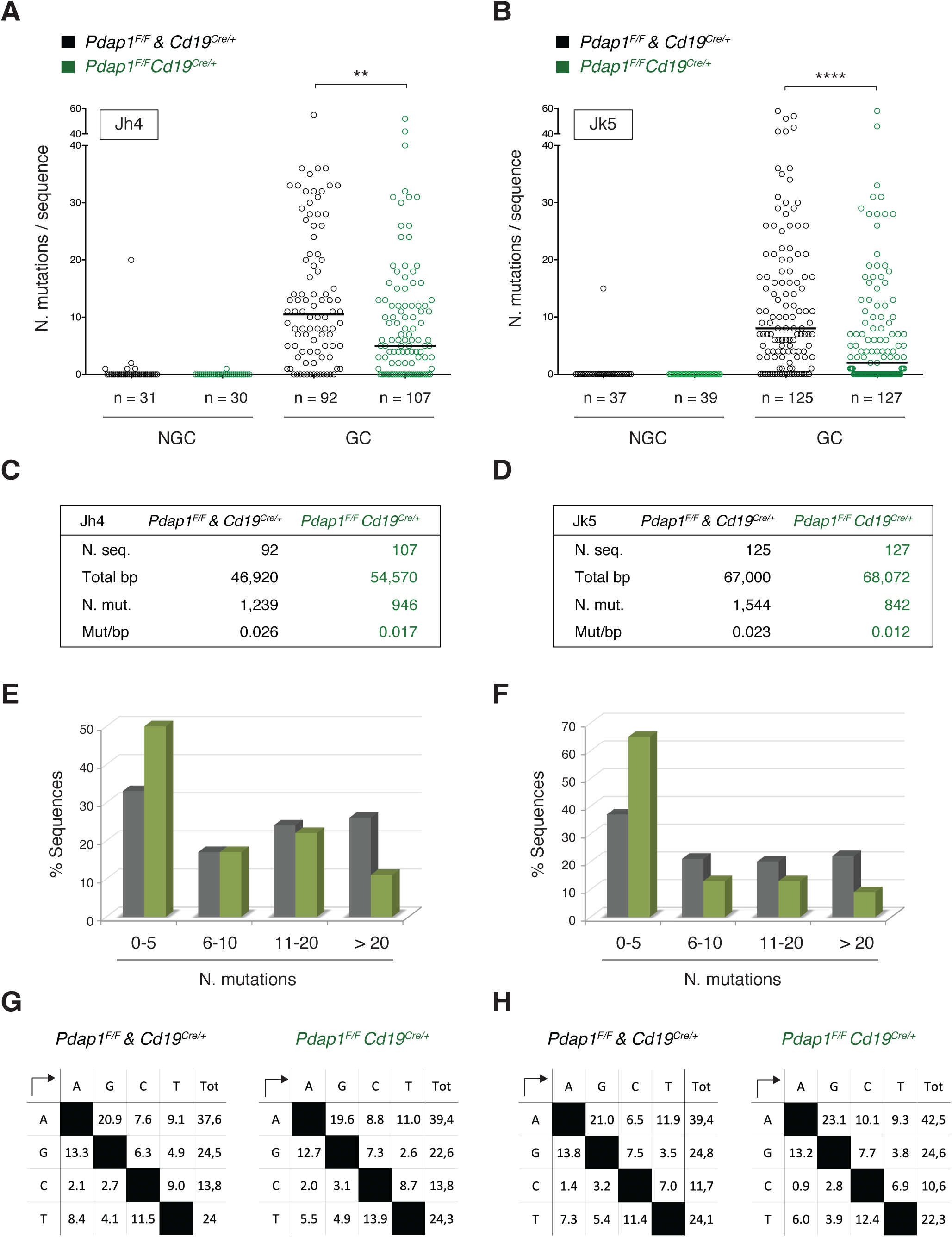
Pdap1 is required for efficient SHM of *Ig* loci. **A and B.** Graphs summarizing number of mutations in 3’ J_H_4 (A) and 3’ J_k_5 (B) regions cloned from sorted Peyer’s patches B cells of aged, unimmunized mice (5 mice per genotype). Each symbol in the graphs indicates a single sequence, and the total number of sequences analysed for each group is indicated below. Mutations were quantified over 510 bp downstream J_H_4 gene segment (A) and 536 bp downstream J_K_5 gene segment (B). NGC: non-germinal center; GC: germinal center. **C and D.** Summary tables listing number of analyzed sequences and total length, number of mutations, and mutation frequency at J_H_4 (C) and J_K_5 (D) introns from panels A and B. **E and F.** Graphs showing the percentage of sequences from panels A and B bearing the indicated mutations in 3’ J_H_4 (E) and 3’ J_k_5 (F). **G and H.** Profiles of nucleotide substitutions at 3’ J_H_4 (G) and 3’ J_k_5 (H) regions. Significance in panels A and B was calculated with the Mann–Whitney U test, and the median is indicated. ** = p ≤ 0.01; **** = p ≤ 0.0001.

Altogether, these findings indicate that ablation of Pdap1 impairs the efficiency of SHM of heavy and light chain *Ig* loci. Therefore, Pdap1 is required to support both antibody diversification reactions occurring in mature B cells, namely CSR and SHM.

## Discussion

In this study, we report the identification of a novel factor required for mature B cell homeostasis and function. Pdap1-deficient B cells develop normally, but display reduced cell viability at the mature stage both *in vitro* and *in vivo*. Furthermore, they fail to efficiently induce AID expression, and cannot support physiological levels of CSR and SHM.

Pdap1 deficiency in mature B cells phenocopies key aspects of the cellular response to chronic stress. Stress-induced eIF2*α* phosphorylation drives the inhibition of global protein synthesis, while allowing the preferential translation of stress responsive genes like Atf4 (Harding et al., 2000; Hinnebusch, 2000; Scheuner et al., 2001; Lu et al., 2004). The transcriptional program induced by Atf4 comprises cytoprotective genes as well as the upregulation of biosynthetic pathways to reprogram cellular metabolism and aid homeostasis recovery. This anabolic program encompasses metabolic pathways of amino acid biosynthesis and the expression of amino acid transporter and aminoacyl-tRNA synthetases genes, and supports recovery of translation under conditions of acute stress (Harding et al., 2003; Rendleman et al., 2018; Quirós et al., 2017; Krokowski et al., 2013; Han et al., 2013). However, during chronic stress, Atf4-dependent increase in protein synthesis leads to proteotoxicity, oxidative stress and cell death (Han et al., 2013; Krokowski et al., 2013). Pdap1-deficient B cells express Atf4 and exhibit the same transcriptional signature of Atf4-mediated translational recovery (Fig. 4, S5, and Table S1). Furthermore, the expression of this anabolic program correlates with increased apoptosis of *Pdap1^F/F^Cd19^Cre/+^* B cells. Therefore, our data is consistent with a model in which ablation of Pdap1 in activated B cells leads to cell death *via* induction of an Atf4-dependent chronic stress response. In addition, resting B cells from Pdap1-deficient mice displayed phosphorylation of Perk and its downstream target eIF2*α* as well as Atf4 expression, thus indicating constitutive activation of the Perk-mediated ISR pathway. This result is in agreement with the reduced number of resting B cells isolated from spleens of *Pdap1^F/F^Cd19^Cre/+^* (Fig. S1E), and indicates that Pdap1-deficiency is detrimental also in naïve B cells. Therefore, Pdap1 is required to protect B lymphocytes from stress-induced cell death under both resting and activated conditions.

Interestingly, while Atf4 expression increased steadily over time after activation in Pdap1-deficient B cells, phosphorylation of Perk and eIF2*α* was no longer detectable. This observation has two important implications. First, the upregulation of Atf4 expression in Pdap1-ablated cells is uncoupled from Perk-eIF2*α* phosphorylation following activation. This result is in agreement with the identification of a phospho-eIF2*α*-independent regulation of Atf4 expression observed during conditions of prolonged cellular stress (Guan et al., 2014). This regulation is likely under the control of mTORC1, which is activated in B cells after stimulation (Powell et al., 2012; Limon and Fruman, 2012; Iwata et al., 2017), and has been shown to promote Atf4 translation in a phospho-eIF2*α*-independent manner in other cellular contexts (Park et al., 2017; Ben-Sahra et al., 2016). Second, the dephosphorylation of Perk and eIF2*α* occurs at early time points after activation. This observation implies the existence of an active mechanism operating immediately upon B cell activation that physiologically inhibits the Perk-eIF2*α* signaling cascade. In line with this point, although mature B cells retain the potential to activate all three arms of the UPR in response to pharmacological triggers of the pathway, the Perk branch is suppressed during normal differentiation of activated B cells into plasma cells (Ma et al., 2010; Gass et al., 2008; Zhang et al., 2005). It is interesting to notice that non-physiological Perk-eIF2*α*-Atf4 signaling after B cell activation blocks the formation of plasma cells (Zhu et al., 2019). In *Pdap1^F/F^Cd19^Cre/+^* splenocytes, the Perk-eIF2*α*-Atf4 pathway is already active under resting conditions, and the increased loss of mitochondrial membrane potential and apoptosis were observed at early, but not late, stages after activation. The earlier kinetics of Atf4 induction in our experimental conditions would explain why Pdap1-deficiency leads to cell death of naïve and early activated B cells but it does not affect the plasma cell compartment.

The activation of the Atf4-dependent stress response in *Pdap1^F/F^Cd19^Cre/+^* B cells was accompanied by downregulation of key factors driving antibody diversification, including AID (Muramatsu et al., 2000; Revy et al., 2000) and the transcription factor Batf, which has been shown to directly control AID expression and other aspects of the CSR program (Ise et al., 2011). Defective induction of *Aicda* transcription in activated Pdap1-deficient B cells was common to all stimulation conditions (Fig. 3A and 4C), and likely represents the dominant cause for the reduced CSR and SHM efficiency. *Aicda* is an haploinsufficient gene, and the efficiency of CSR is directly proportional to AID levels. Mice heterozygous for a null *Aicda* allele display defects in both CSR and SHM (McBride et al., 2008; Sernández et al., 2008; Takizawa et al., 2008). Analogously, Pdap1 deficiency reduced *Aicda* expression to approximately half of wild-type levels in activated B cells, and is associated with a similar reduction in the efficiency of both antibody diversification reactions. Furthermore, overexpression of AID in LPS-BAFF-TGFβ-stimulated splenocytes, which did not exhibit any additional defect in GLT, rescued the IgG2b CSR defect of *Pdap1^F/F^Cd19^Cre/+^* B cells to a considerable extent. Although we cannot exclude the possibility that Pdap1 might regulate the expression of AID independently from its protective role against chronic ISR activation, it is intriguing to speculate that the sustained induction of the ISR in *Pdap1^F/F^Cd19^Cre/+^* B cells might actively interfere with the CSR and SHM programs. In this regard, limiting AID expression under conditions of cellular stress might represent a mechanism to alleviate the additional burden imposed by AID-induced genotoxic stress on activated B cells.

## Acknowledgements

We thank all members of the Di Virgilio lab for their feedback and discussion; V. Coralluzzo (AG Di Virgilio, MDC, Berlin) for support with genotyping; Y. Dramaretska (AG Gargiulo, MDC) for helpful tips on the RNA-Seq protocol; the MDC Transgenics platform and Dr. R. Kühn for the generation of the *Pdap1^F^* mouse allele; the MDC FACS Core Facility and Dr. HP. Rahn for assistance with cell sorting; K. Rajewsky (MDC) for feedback and discussion; and N. Zampieri (MDC) for critical reading of the manuscript and advices. We are particularly grateful to M. Caganova (AG K. Rajewsky, MDC) for sharing her expertise and reagents for the ISR and UPR pathway analyses. This work was supported by ERC grant 638897 (to M.D.V.), the Helmholtz-Gemeinschaft Zukunftsthema “Immunology and Inflammation” ZT-0027 (to M.D.V.), and the Deutsche Krebshilfe grant 70112800 (to M.J.). M.D.V. is a Helmholtz Young Investigators Group leader (Helmholtz Association).

## Author contributions

M.D.V. and V.D.B. conceived the project idea and designed the experiments; M.D.V., V.D.B., and M.B.L. analyzed and interpreted the data; V.D.B., M.B.L., W.W., S.B., D.S., A.R., M.D., and L.K. performed the experiments; R.A. performed all RNA-seq data analyses, and R.G. contributed to design the list of gRNAs used for the initial loss-of-CSR screen; M.J., A.A., and M.D.V. supervised the experiments and/or analyses performed in their respective groups; M.D.V. obtained the funding for the project and wrote the manuscript.

## Supplementary Figure Legends

**Figure S1.**
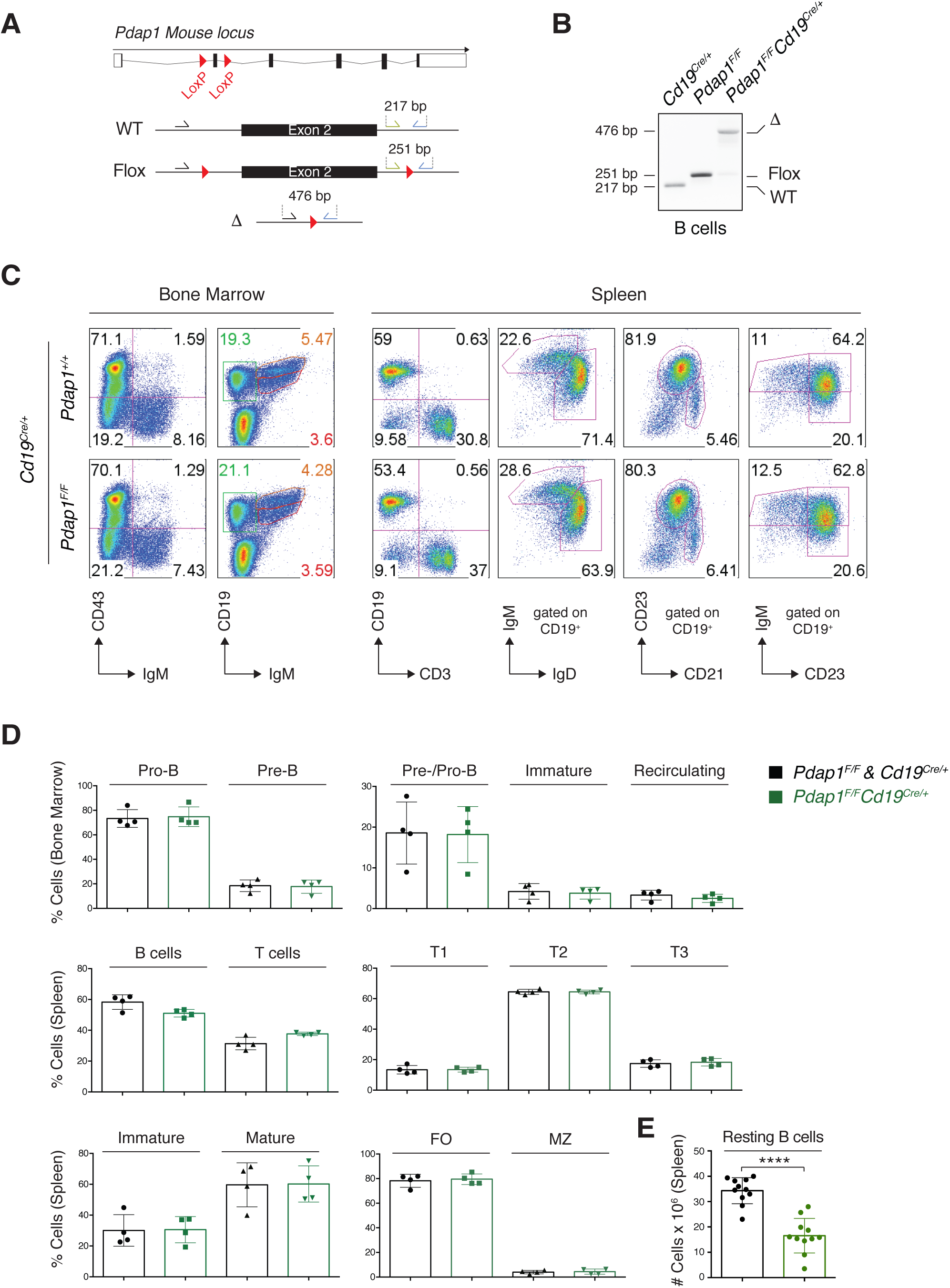
Pdap1 is largely dispensable for B cell development. **A.** Schematic representation of *Pdap1* locus and genotyping strategy. WT: wildtype *Pdap1* allele; Flox: conditional *Pdap1* allele; Δ: null *Pdap1* allele. **B.** Electrophoretic analysis of *Pdap1* allele status in resting splenic B cells isolated from mice of the indicated genotype. **C.** Representative flow cytometry analysis of lymphoid tissues from control and *Pdap1^F/F^Cd19^Cre/+^* mice. **D.** Summary graphs for 4 mice per genotype. **E.** Number of resting B cells isolated from spleens of the indicated genotype. Each dot represents a different mouse. Significance in panels D and E was calculated with the Mann–Whitney U test, and error bars represent SD. No difference among groups was significant for graphs in panel D. **** = p ≤ 0.0001.

**Figure S2.**
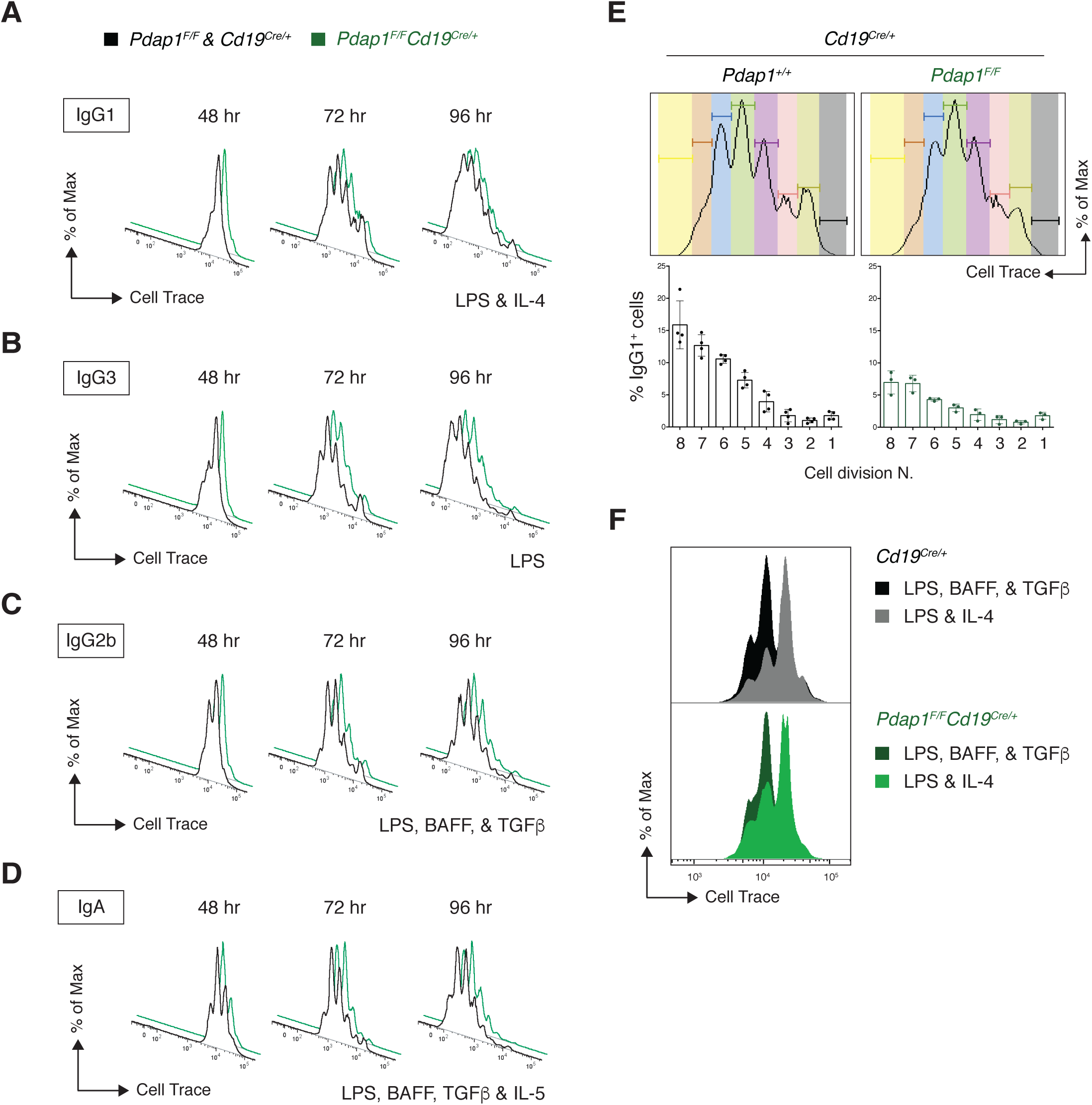
Pdap1 is dispensable for B cell proliferation. **A-D.** Proliferation analysis by CellTrace Violet dilution of primary cultures of *Cd19^Cre/+^* and *Pdap1^F/F^Cd19^Cre/+^* B lymphocytes stimulated with LPS-IL-4 (A), LPS only (B), LPS-BAFF-TGFβ (C), or LPS-BAFF-TGFβ-IL-5 (D). Data are representative of at least 2 mice per genotype. **E.** Graph showing percentage of IgG1^+^ cells per cell division in primary cultures of *Cd19^Cre/+^* and *Pdap1^F/F^Cd19^Cre/+^* splenocytes stimulated with LPS and IL-4 for 72 h. Graph summarizes at least 3 mice per genotype, and error bars represent SD. Representative cell division plots as measured by CellTrace Violet dilution are shown on top. **F.** Representative overlay of CellTrace Violet proliferation tracks of LPS-IL-4 and LPS-BAFF-TGFβ cultures at 48 h post-activation.

**Figure S3.**
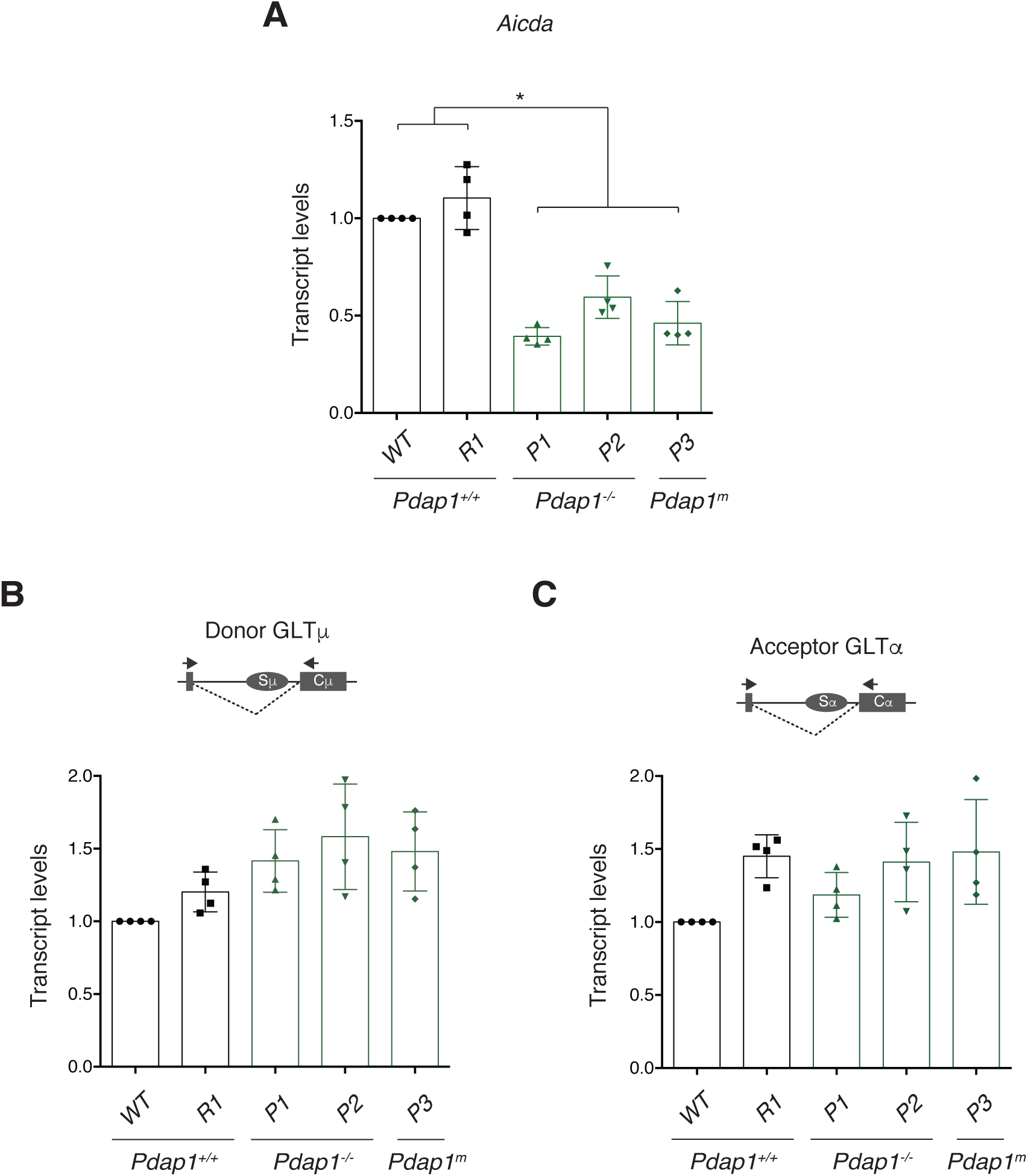
*Aicda* mRNA levels are reduced in Pdap1-deficient CH12 cell lines. **A.** qPCR analysis for *Aicda* mRNA in activated CH12 cell lines of the indicated genotypes. Graph summarizes four independent experiments. Parental WT CH12 cells within each experiment were assigned an arbitrary value of 1. **B and C.** qPCR analysis for Igμ (B) and Ig*α* (C) GLT levels in activated CH12 cell lines of the indicated genotypes. The schematic representations on top of each graph indicate the location of primers employed to analyze post-spliced germline transcripts. Graphs summarize four independent experiments. Parental WT CH12 cells within each experiment were assigned an arbitrary value of 1. Significance in panel A was calculated with the Mann–Whitney U test. Error bars represent SD. * = p ≤ 0.05.

**Figure S4.**
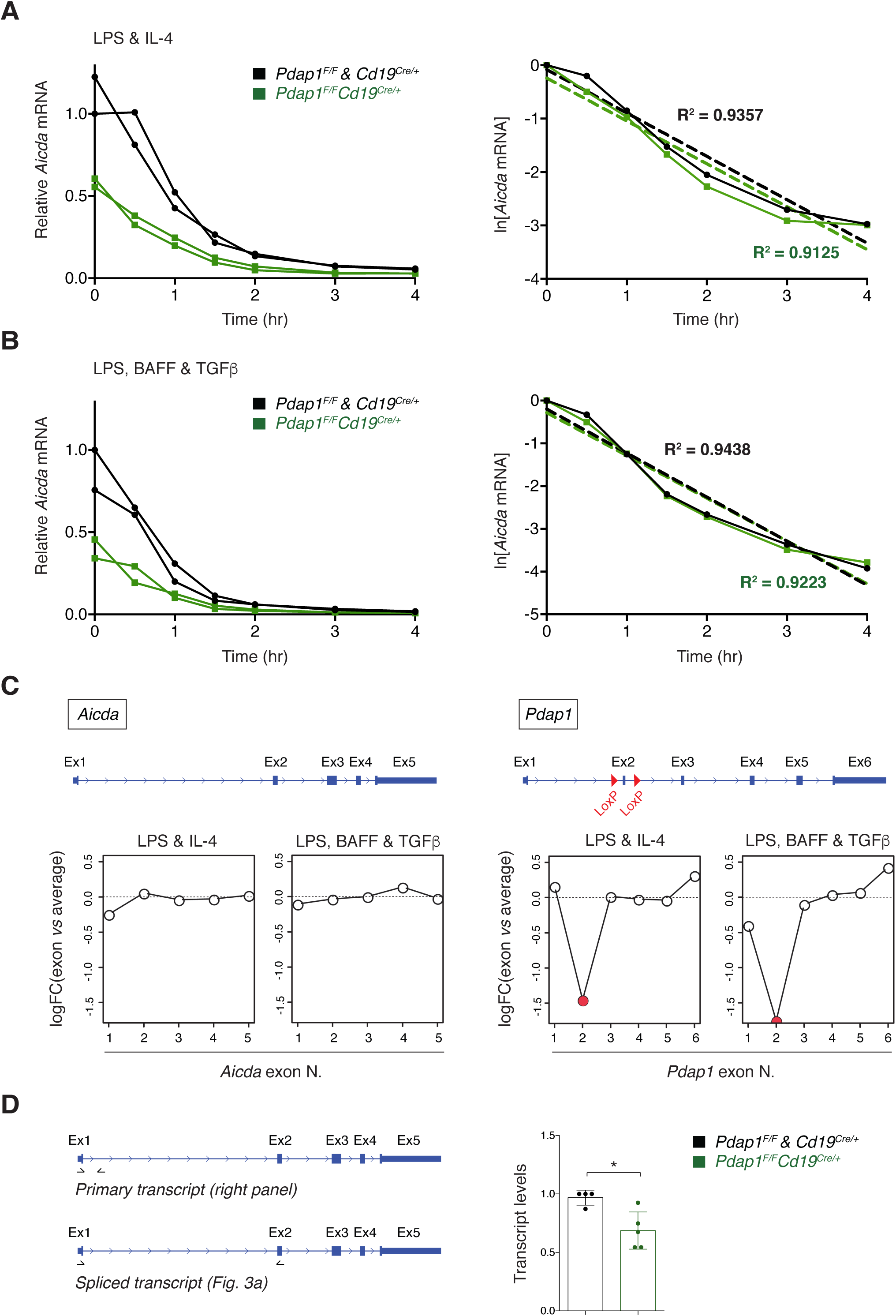
Pdap1 is dispensable for *Aicda* mRNA stability and splicing. **A-B.** Left: qPCR analysis for *Aicda* mRNA from B cells of the indicated genotypes stimulated with LPS and IL-4 (A) or LPS, BAFF and TGFβ (B) for 48 h to induce CSR to IgG1 or IgG2b, respectively, and treated with Actinomycin-D for the indicated time. Graph summarizes the mean of two independent qPCR measurements per mouse for 2 mice per genotype. Right: Linear regression analysis of same data, shown as ln[RNA] *versus* time of Actinomycin-D treatment. **C.** Analysis of exon usage for *Aicda* (left) and *Pdap1* (right) genes. *Pdap1* splicing data is provided as a positive control for the spicing analysis. A schematic representation of each gene is provided above. **D.** Left: Schematic representation of *Aicda* gene with location of primers employed to analyze *Aicda* primary and spliced transcripts. Right: qPCR analysis of *Aicda* primary transcripts in B cells 48 h after activation with LPS. Graphs summarize at least 4 mice per genotype. Results of qPCR analysis of *Aicda* spliced transcripts are shown in Fig. 3A. One *Cd19^Cre/+^* mouse within each experiment was assigned an arbitrary value of 1. Significance in panel D was calculated with the Mann–Whitney U test, and error bars represent SD. * = p ≤ 0.05.

**Figure S5.**
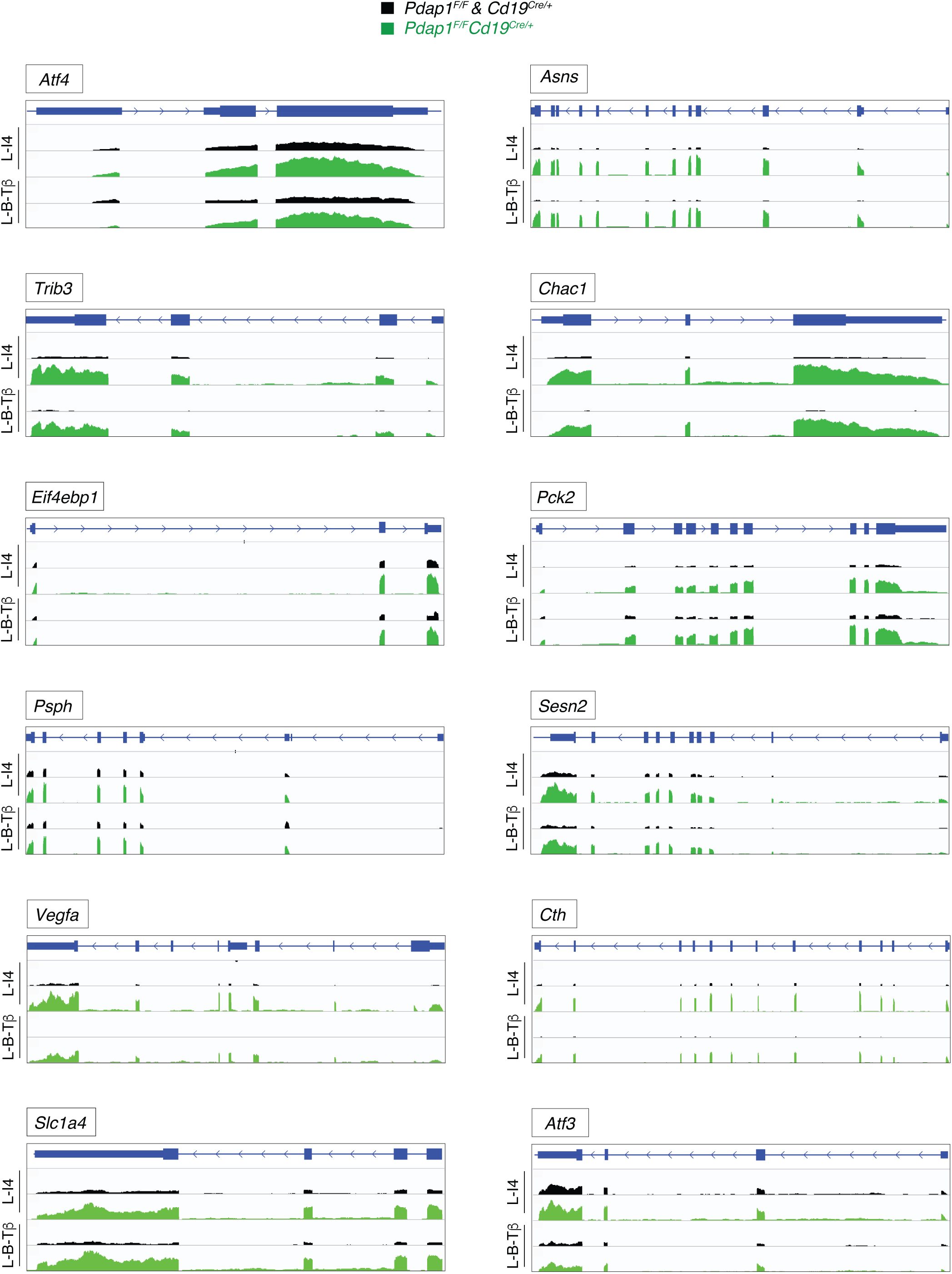
Increased expression of stress response genes in Pdap1-deficient B cells. Coverage of stress response genes in B cells of the indicated genotypes 48 h after activation with LPS and IL-4 (L-I4) or LPS, BAFF, and TGFβ (L-B-Tβ), as measured by RNA-Seq analysis. Twelve selected genes are shown.

**Table S1.** Pairwise comparison of differentially regulated genes in control *versus Pdap1^F/F^Cd19^Cre/+^* splenocytes.

**Table S2.**
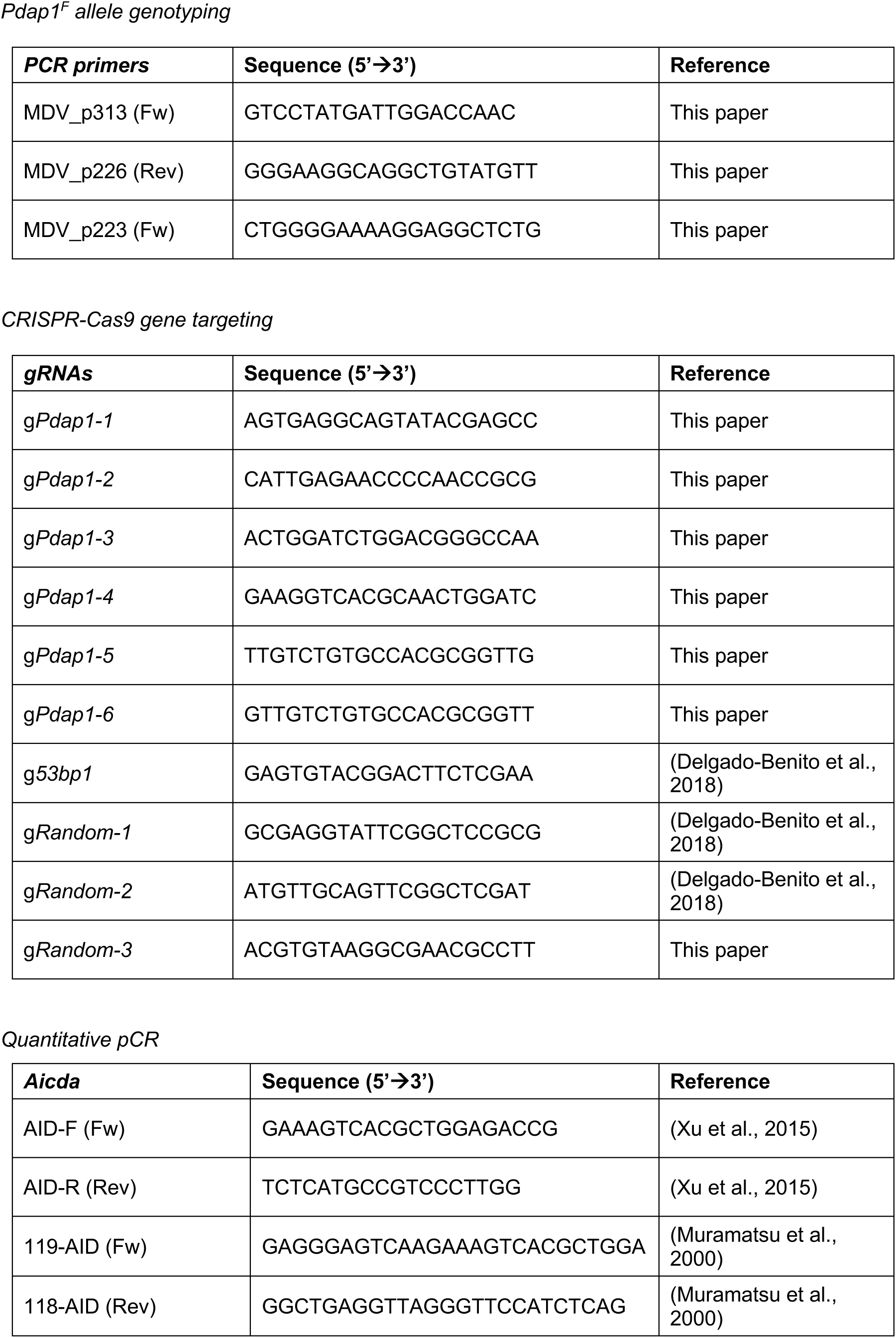
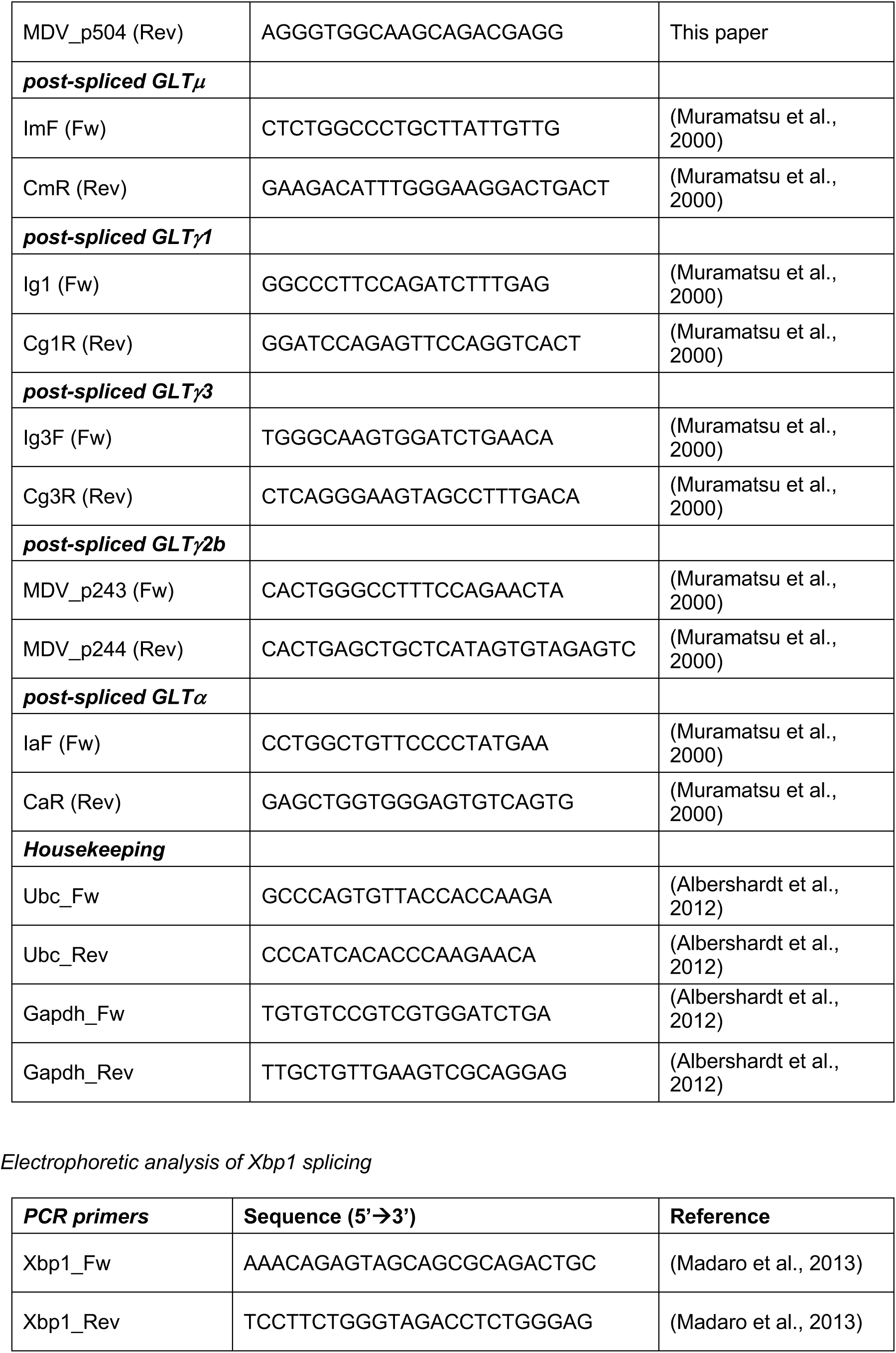
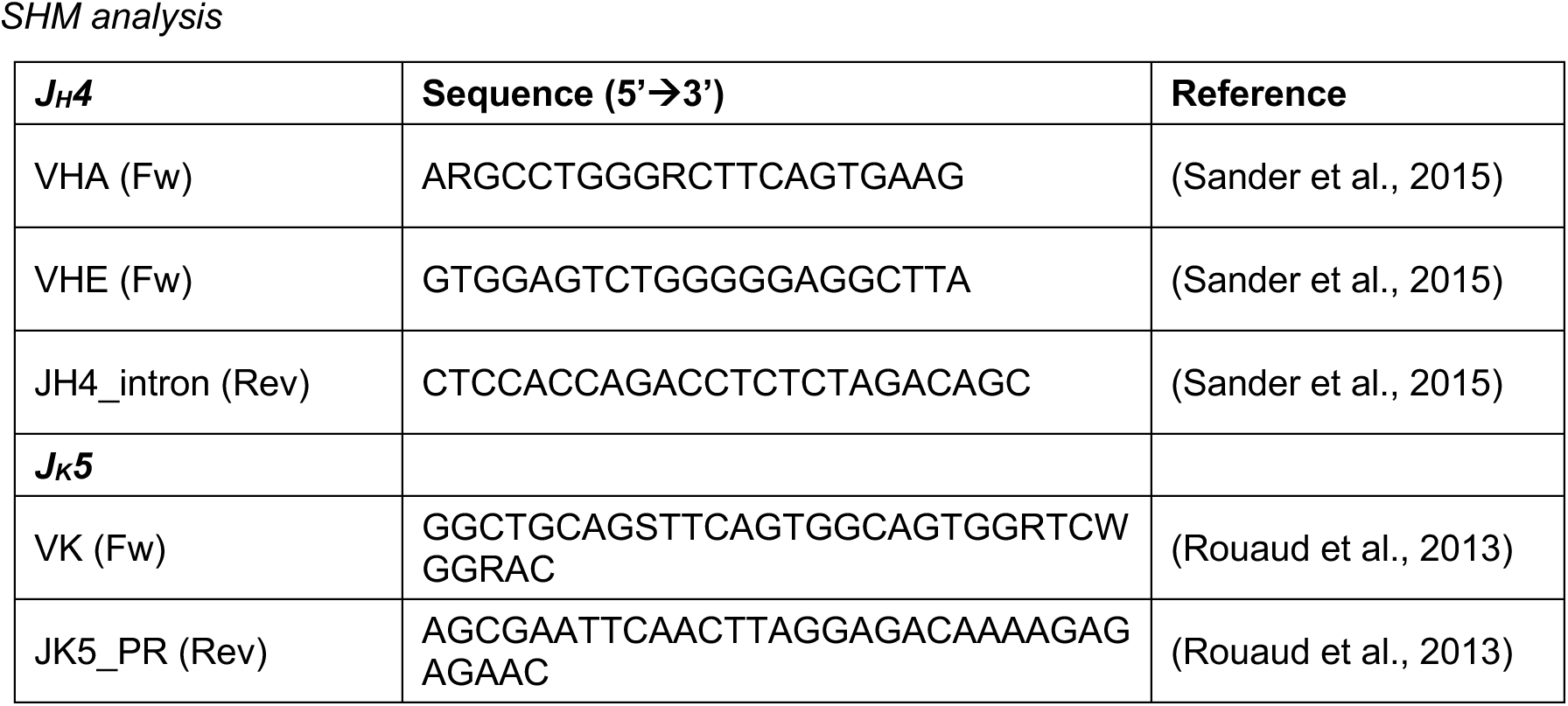
List of oligonucleotides used in this study.

## Methods

### Mice strains

*Cd19^Cre^* (*Cd19^tm1(cre)Cgn^*, (Rickert et al., 1997)) and *Aicda^−/−^* (*Aicda^tm1Hon/tm1Hon^*, (Muramatsu et al., 2000)) mice were previously described. The conditional *Pdap1^F^* allele bears LoxP sites flanking Exon 2 (ENSMUST00000031627.8), and was generated by CRISPR/Cas9-mediated knock-in microinjection of single cell embryos on a C57BL/6N strain background (MDC Transgenics platform). Germline transmission was confirmed and positive pups were bred with *Cd19^Cre^* mice to generate *Pdap1^F/F^Cd19^Cre/+^* mice. Mice were maintained in a specific pathogen-free (SPF) barrier facility under standardized conditions (20+/-2 °C temperature; 55%±15% humidity) on a 12 h light/12 h dark cycle. Mice of both genders 7 to 27 weeks old were used in age-matched groups for the experiments. All experiments were performed in compliance with the European Union (EU) directive 2010/63/EU, and in agreement with Landesamt für Gesundheit und Soziales directives (LAGeSo, Berlin, Germany). Primers used for genotyping the *Pdap1^F^* allele are listed in Table S2.

### Primary cell cultures

B lymphocytes were isolated from mouse spleens using anti-CD43 MicroBeads (Miltenyi Biotec), and grown in RPMI 1640 medium (Life Technologies) supplemented with 10% fetal bovine serum (FBS), 10 mM HEPES (Life Technologies), 1 mM Sodium Pyruvate (Life Technologies), 1X Antibiotic Antimycotic (Life Technologies), 2 mM L-Glutamine (Life Technologies), and 1X 2-Mercaptoethanol (Life Technologies) at 37 °C and 5% CO2 levels.

### Cell lines

The cell lines employed for this study are: CH12 (CH12F3, murine, (Nakamura et al., 1996)), WT (R1 and R2) and Pdap1-deficient (P1, P2, and P3) CH12 clonal derivatives (murine, this paper), and BOSC23 (human, (Pear et al., 1993)). CH12 cells were grown in RPMI 1640 medium supplemented with 10% fetal bovine serum (FBS), 10 mM HEPES, 1 mM Sodium Pyruvate, 1X Antibiotic Antimycotic, 2 mM L-Glutamine, and 1X 2-Mercaptoethanol at 37 °C and 5% CO2 levels. BOSC23 cells were cultured in DMEM medium (Life Technologies) supplemented with 10% FBS, 2 mM L-Glutamine, and Penicillin-Streptomycin (Life Technologies) at 37 °C and 5% CO2 levels.

### CSR assay

CH12 cells were stimulated to undergo CSR to IgA by treatment with 1-5 μg/mL *α*CD40 (BioLegend), 5 ng/ml TGFβ (R&D Systems) and 5 ng/ml of mouse recombinant IL-4 for 48 h. B lymphocytes were stimulated to undergo class switching with 5 μg/ml LPS (Sigma-Aldrich) and 5 ng/ml of mouse recombinant IL-4 (Sigma-Aldrich) for CSR to IgG1; 5 μg/ml LPS only for CSR to IgG3; 5 μg/ml LPS, 10 ng/ml BAFF (PeproTech) and 2 ng/ml TGFβ for CSR to IgG2b; or 5 μg/ml LPS, 10 ng/ml BAFF, 2 ng/ml TGFβ and 1.5 ng/ml recombinant murine IL-5 (PeproTech) for CSR to IgA. For class switching analysis, cell suspensions were stained with fluorochrome-conjugated anti-IgG1, anti-IgG3 (BD-Biosciences), anti-IgG2b (BioLegend), or anti-IgA (Southern Biotech).

### Retroviral infection

The pMX-AID-ER-IRES-GFP retroviral vector was a kind gift from Qiao Wang (The Rockefeller University, New York). Splenocytes infections were performed as it follows. The 293T derivative cell line BOSC23 was transfected with pCL-Eco and pMX-IRES-GFP or pMX-AID-ER-IRES-GFP retroviral vectors using FuGENE® HD Transfection Reagent (Promega) to generate viral particle-containing supernatants. B-cells were activated with 5 μg/ml LPS and 5 ng/ml IL-4 (IgG1) or 5 μg/ml LPS, 10 ng/ml BAFF and 2 ng/ml TGFβ (IgG2b), and transduced twice with the viral supernatant, one and two days after isolation. 4-hydroxytamoxifen (4-HT) was added 72 h post-activation at a final concentration of 0.05 μM, and CSR was assessed 24 h after on the GFP^+^-gated populations.

### B cell development and differentiation analyses

For analysis of B cell development and differentiation, spleen and bone marrow cell suspensions were incubated with ACK lysis buffer (ThermoFisher) for erythrocytes depletion. Subsequently, cells were blocked with TruStain fcX (BioLegend) for 10 min at 4 °C and labeled with fluorochrome conjugated antibodies to determine the surface expression of CD43, IgM, CD21/CD35, IgD (BD Biosciences) and CD23, CD3 and CD19 (BioLegend) in PBS supplemented with 3 % FBS (PBS/FBS) for 20 min at 4 °C. Cells were then washed, resuspended in PBS/FBS and analyzed.

For analysis of plasma cell differentiation *in vitro*, splenic B cells were isolated by immunomagnetic depletion of CD43^+^ cells, and cultured at a density of 10^6^ cells/ml in the presence of either 20 µg/ml LPS (Sigma, #L2880-10MG) and 25 ng/ml IL-4, or 20 µg/ml LPS, 10 ng/ml BAFF, and 2 ng/ml TGFβ. To determine the percentage of plasmablasts, cells were harvested after 96 hours, blocked with TruStain fcX for 10 min at 4 °C, and subsequently stained for surface expression of B220 (BioLegend, #103245) and CD138 (BioLegend, #142506) in MACS buffer (PBS supplemented with 0.5 % BSA and 2 mM EDTA) for 20 min at 4 °C. Cells were then resuspended in FACS buffer (PBS supplemented with 3 % FCS and 1 mM EDTA) containing 1 µg/ml propidium iodide and analyzed.

For analysis of the plasma cell compartment *in vivo*, 9, 14 and 25 weeks old mice were sacrificed to isolate the spleen and hind legs. Total splenocytes and bone marrow cells were isolated by smashing the respective organs and erythrocytes were depleted by incubation with Gey’s ABC solution for 3 min on ice. For surface staining, 5×10^6^ cells were first blocked with TruStain fcX for 10 min at 4 °C and then stained for the following markers for 20 min at 4 °C in MACS buffer: CD138, TACI (BD Pharmingen, #558410), CD93 (BioLegend, #136506), CD19 (BioLegend) and B220. Cells were resuspended in FACS buffer containing 1 µg/mL propidium iodide and analyzed.

For assessment of germinal center B cells and CSR *in vivo*, cell suspensions derived from Peyer’s Patches were blocked with TruStain fcX for 10 min at 4 °C. Subsequently, cells were stained with fluorochrome-conjugated anti-CD19, anti-B220/CD45R, anti-CD38 (BioLegend) anti-Fas/CD95 and anti-IgA (BD Biosciences) for 20 min at 4 °C in PBS/FBS. Cells were washed, resuspended in PBS/FBS and analyzed.

All samples were acquired on a LSRFortessa cell analyzer (BD-Biosciences).

### Cell proliferation and apoptosis analysis

For cell proliferation analysis by cell tracking dye dilution, primary B cells were pulsed with 5 μM CellTrace Violet (Thermofisher) for 10 min at 37 °C. Apoptosis analysis was performed by using CaspGLOW™ Fluorescein Active Caspase Staining Kit (BioVision) according to the manufacturer’s instructions. Samples were acquired on a LSRFortessa cell analyzer (BD-Biosciences). Mitochondrial mass and membrane potential were measured *via* staining with MitoTracker Green and DeepRed (ThermoFisher), respectively, according to the manufacturer’s instructions. Samples were acquired on a LSRFortessa cell analyzer (BD-Biosciences).

### CRISPR-Cas9 gene targeting

For the loss-of-CSR assay in bulk CH12 cultures, 3 gRNAs against *Pdap1* gene were cloned into the U6 cassette of a variant of the original pX330 plasmid (pX330-U6-Chimeric_BB-CBh-hSpCas9, Addgene #42230) modified to express Cas9^WT^-T2A-GFP (kind gift from Van Trung Chu, MDC) (g*Pdap1-1*/*3*), and 3 gRNA pairs were cloned into tandem U6 cassettes in a mutated version of pX330 expressing Cas9^D10A^ (*Nickase-a/c*). CH12 cells were transfected with the Cas9-gRNAs expressing constructs (either individually or in a pooled format) *via* electroporation with Neon Transfection System (Thermo Fisher Scientific), sorted for GFP-positive cells after 40 h, and left to recover for 72 h before activation for CSR analysis. For the generation of Pdap1-deficient CH12 clonal derivatives, the *Nickases-a*/*c* constructs were individually electroporated into CH12. Single GFP-positive cells were sorted in 96-well plates after 40 h, and clones were allowed to grow for 12 days before CSR analysis in 96-well format and expansion of selected clones. Controls for gRNA-nucleofected CH12 were cells nucleofected either with empty vector or gRNAs against random sequences not present in the mouse genome (single random gRNA-Cas9^WT^ as controls for g*Pdap1-1*/*3*, and random gRNAs pairs-Cas9^D10A^ as controls for *Nickase-a/c*). Selected clones R1/R2 and P1/3 were validated at the level of genomic scar and protein expression. The sequences of the gRNAs employed in these studies (this paper; (Delgado-Benito et al., 2018)) are listed in Table S2.

### Western blot analysis

Western blot analysis of protein levels was performed on whole cell lysates prepared by lysis in RIPA buffer (Sigma) supplemented with Complete EDTA free proteinase inhibitors (Roche). Pierce Phosphatase Inhibitor Mini Tablets (ThermoFisher) were added to the lysis buffer for the analysis of phosphorylated Perk and eIF2*α*. The antibodies used for WB analysis are: anti-Pdap1 (Sigma), anti-Tubulin (Abcam), anti-βActin (Sigma), anti-AID (mAID-2, ThermoFisher), anti-phospho-Perk (pThr980, ThermoFisher), anti-Perk (Cell Signaling Technology), anti-phospho-eIF2*α* (pSer51, Cell Signaling Technology), anti-eIF2*α* (Santa Cruz), anti-Atf4 (Cell Signaling Technology), and anti-Xbp1-s (Cell Signaling Technology).

### RT- and quantitative PCR

mRNA levels for AID and post-spliced germline transcripts were measured as it follows. Total RNA was extracted from splenocytes cultures 48 h after activation using TRIzol (Invitrogen) according to manufacturer’s instructions, and retro-transcribed with SuperScript VILO cDNA Synthesis Kit (Invitrogen). Genomic DNA was removed by RapidOut DNA Removal Kit (Thermo Scientific). Transcripts were amplified using StepOnePlus Real-Time PCR System (Applied Biosystems) with Luna Universal qPCR Mastermix (NEB). For analysis of *Aicda* mRNA decay, Actinomycin D was added to the culture medium 48 h post-activation at a final concentration of 10 μg/ml for 0.5, 1, 1.5, 2, 3, and 4 hr. qPCR analyses were normalized to *Ubc* or *Gapdh* (*Aicda* mRNA and GLTs) or *Gapdh* (*Aicda* mRNA decay). Primers used for qPCR (this paper; (Xu et al., 2015; Muramatsu et al., 2000; Albershardt et al., 2012)) are listed in Table S2.

For electrophoretic analysis of *Xbp1* splicing, *Xbp1* transcripts were amplified using the primers listed in Table S2 (Madaro et al., 2013), and either mock-digested or digested with *PstI* enzyme (NEB) for 30 min at 37 °C. The intact spliced (Xbp1-s) transcript RT-PCR product and the unspliced (Xbp1-u) transcript digestion products were visualized on 2% agarose gel.

### RNA-Seq

RNA-Seq analysis was performed on three mice per genotype. Splenocytes were cultured in LPS and IL-4 (IgG1) or LPS, BAFF and TGFβ (IgG2b) for 48 h. Cells were collected by centrifugation and RNA was extracted with TRIzol (Invitrogen) according to manufacturer’s instructions. Ribosomal RNA was depleted using Ribo-Zero Gold rRNA Removal Kit (Illumina). Libraries were prepared with TruSeq Stranded Total RNA Library Prep Kit Gold (Illumina), and run in one lane on a flow cell of HiSeq 4000 (Illumina).

The RNA-seq data was analysed using the pigx-rnaseq pipeline (Wurmus et al., 2018). STAR (Dobin et al., 2013) mapped the reads to the GRCm38 assembly for mouse by Ensembl, and HTSeq (Anders et al., 2015) was used to count the transcript abundance. Differential expression analysis was done using the DESeq2 (Love et al., 2014) package for R, which uses the Wald test for significance.

For splicing analysis of *Aicda* and *Pdap1* genes, we used the edgeR (Robinson et al., 2010) package for R to determine differentially expressed exons. The annotation was provided by a filtered version of the GRCm38 gene annotation, which contained unique, Havana annotated exons.

### SHM analysis

Single-cell suspensions from Peyer’s patches of 24-27 weeks old mice were first incubated with TruStain fcX, and then labeled with B220-FITC- (BioLegend), CD19-APC/Pacific Blue (BioLegend), CD38-Alexa700- (Thermo Scientific), and CD95/Fas- PE- (BD-Biosciences) conjugated antibodies. Non-germinal center (CD38^+^ Fas^−^) and germinal center B cells (CD38^−^ Fas^+^) were sorted on an Aria BD sorter. Genomic DNA was extracted and the 5’ portions of J_H_4 (*Igh*) and J_K_5 (*Igk*) introns were amplified by PCR using Phusion High-Fidelity DNA Polymerase (Thermo Scientific). The 800 bp J_H_4 and 700 bp J_K_5 PCR products were gel extracted, cloned into a pCR2.1 vector using the TOPO TA Cloning Kit (Invitrogen) and sequenced.

Mutations were quantified over 510 bp downstream J_H_4 and 536 bp downstream J_K_5 gene segments. Primers used for SHM analysis (Rouaud et al., 2013; Sander et al., 2015) are listed in Table S2.

### Statistical analysis

The statistical significance of differences between groups/datasets was determined by the Mann–Whitney U test for all data presented in this study, with the following exceptions. For the RNA-Seq analysis, the Wald test of the DESeq2 package for R was used, and we considered genes with an FDR < 0.05 to be significantly differentially expressed. Statistical details of experiments can be found in the figure legends.

### Data availability

The deep-sequencing data reported in this paper (RNA-Seq) has been deposited in the GEO repository under accession number GSE141876.

